# MYC regulates a pan-cancer network of co-expressed oncogenic splicing factors

**DOI:** 10.1101/2021.11.24.469558

**Authors:** Laura Urbanski, Mattia Brugiolo, SungHee Park, Brittany Angarola, Nathan K. Leclair, Phil Palmer, Sangram Keshari Sahu, Olga Anczuków

## Abstract

MYC is dysregulated in >50% of cancers, but direct targeting of MYC has been clinically unsuccessful. Targeting downstream MYC effector pathways represents an attractive alternative. MYC regulates alternative mRNA splicing, a hallmark of cancer, but the mechanistic links between MYC and the splicing machinery remain underexplored. Here, we identify a network of splicing factors (SFs) co-expressed as SF-modules in MYC-active breast tumors. Of these, one is a pan-cancer SF-module, correlating with MYC-activity across 33 tumor types. In mammary cell models, MYC activation leads to co-upregulation of pan-cancer module SFs and to changes in >4,000 splicing events. In breast cancer organoids, co-overexpression of the pan-cancer SF-module is sufficient to induce splicing events that are also MYC-regulated in patient tumors and to increase organoid size and invasiveness, while its knockdown decreases organoid size. Finally, we uncover a pan-cancer splicing signature of MYC activity which correlates with survival in multiple tumor types. Our findings provide insight into the mechanisms and function of MYC-regulated splicing and for the development of therapeutics for MYC-driven tumors.

## INTRODUCTION

Alternative RNA splicing is a key step in gene expression regulation, contributing to transcriptomic and proteomic diversity by controlling which exons are joined together to produce distinct transcript isoforms. Disruption of splicing in cancer, through mutation and/or altered expression of splicing regulatory factors, impacts the hallmarks of cancer (Bonnal et al., 2020). Mutations in spliceosome components are common in hematological malignancies, whereas solid tumors, including breast, often exhibit changes in copy number and/or expression of splicing factors (SFs) (Urbanski et al., 2018). SFs bind directly to pre-mRNA targets and regulate splicing in a concentration-dependent manner (Long and Caceres, 2009). Indeed, SF expression changes as low as two-fold, in the absence of mutations, have been implicated in splicing deregulation in cancer (Dvinge et al., 2016; Urbanski et al., 2018). Therefore, defining the mechanisms by which SF expression and function are regulated in normal and cancer cells is crucial for understanding SF-mediated transformation and for the development of therapies targeting the splicing machinery (Bonnal et al., 2012; Dvinge et al., 2016)

At the transcriptional level, several SFs are known to be directly regulated by the oncogenic transcription factor MYC through MYC binding sites in their promoter regions (Das et al., 2012; David et al., 2010; Koh et al., 2015; Park et al., 2019). Dysregulation of MYC occurs in more than 50% of all tumors and is associated with poor clinical outcome (Beroukhim et al., 2010; Chen et al., 2018; Gabay et al., 2014). Therapeutic targeting of MYC is therefore expected to have a significant clinical impact, but effective direct targeting of MYC in human tumors has proven difficult due to the lack of a binding site for small molecule inhibitors (Soucek et al., 2008; Wang et al., 2005; Whitfield et al., 2017). Alternative approaches, such as modulating *MYC* transcription and translation or MYC protein stability and activity, show preclinical promise, but there are still no FDA-approved MYC-targeting therapeutics (Chen et al., 2018). Elucidating the molecular mechanisms linking MYC with splicing offers new opportunities for understanding the biology of MYC-driven cancers and for targeting the MYC effector pathway.

In breast tumors, *MYC* is frequently upregulated in the aggressive and difficult to treat basal-like and triple-negative breast cancer subtypes (Xu et al., 2010). In breast cancer models, MYC is known to regulate the expression of individual SFs, such as *SRSF1, TRA2β*, and *BUD31,* and MYC-induced upregulation of these SFs is necessary for MYC-driven tumorigenesis and maintenance (Das et al., 2012; Hsu et al., 2015; Park et al., 2019). Although disrupted SF expression is often observed in human breast tumors, the full extent of SF alterations and their consequences are only beginning to be unraveled. Several studies have experimentally demonstrated that altering the expression of a single SF is sufficient to promote breast tumor formation or metastasis (Anczukow et al., 2012; Jia et al., 2010; Karni et al., 2007; Nguyen et al., 2016; Park et al., 2019). However, among the SFs overexpressed in breast tumors, not all are sufficient, alone, to drive oncogenesis in breast cancer models (Park et al., 2019).

Alternative splicing of a given isoform is the result of combined positive and negative regulation by multiple SFs (Smith and Valcarcel, 2000). In addition, oncogene cooperation has been described for transcriptional and epigenetic regulators. Thus, it is likely that several SFs act together to influence tumorigenesis or tumor maintenance, and, indeed, alterations in multiple SFs are often observed within the same tumor. However, most studies to date have investigated the effects of SFs individually (Anczukow et al., 2012; Ho et al., 2021; Hu et al., 2020; Karni et al., 2007; Kim et al., 2015; Park et al., 2019; Xu et al., 2014; Yang et al., 2016). Further, previous studies have focused on a limited number of MYC-dependent tumor types, but have not investigated if MYC-regulated SFs are tumor-specific or shared across tumor types. Therefore, although SF co-regulation has been postulated to drive cancer progression (Koedoot et al., 2019; Wang et al., 2019a), experimental demonstration of whether multiple SFs can function as a coordinated network of downstream MYC effectors, and whether such SFs have synergistic, cooperative, or even antagonistic effects, is lacking.

Here, we develop a classifier to score tumors as MYC-active or -inactive based on expression of known MYC targets. In breast tumors, we define a splicing signature associated with MYC activity and identify >150 MYC-regulated SFs. We find that MYC-regulated SFs are co-expressed as groups, or modules, six of which have a high correlation with MYC activity. Strikingly, one module is preserved across all tumor types in The Cancer Genome Atlas (TCGA) and maintains a strong correlation with MYC activity. We demonstrate that MYC activation controls the expression of this pan-cancer SF-module in non-transformed mammary epithelial cells. Co-overexpression of three SFs from the pan-cancer module, *SRFS2*, *SRSF3*, and *SRSF7*, leads to increased cell invasiveness in breast cancer models and induces splicing events that are also observed in MYC-active patient tumors. Knockdown of any of these three SFs reduces breast cancer organoid size. Finally, we uncover a splicing signature of MYC-active tumors shared across 75% of analyzed tumor types that correlates with patient survival in multiple cancer types.

## RESULTS

### MYC-active human breast tumors display a distinct splicing signature

We first characterized the splicing landscape in breast tumors with high *vs.* low MYC activity using RNA-sequencing (RNA-seq) data from >1,000 breast tumors in TCGA cohort (Cancer Genome Atlas, 2012; Consortium, 2020). We sought to evaluate MYC activity rather than mRNA expression because the MYC protein requires binding partners to regulate transcription of its targets (Conacci-Sorrell et al., 2014), and because post-translational modifications can alter MYC stability (Hann, 2006). *MYC* mRNA expression is significantly higher in adjacent normal tissue compared to breast tumors in TCGA, highlighting that *MYC* mRNA expression is not an adequate proxy for the function of MYC in tumors (Figure S1A). We therefore adapted a rank-based scoring method (Jung et al., 2017) to define the level of MYC activity across breast samples (Table S1A), where each sample was assigned a MYC activity score based on expression of 200 known MYC target genes in the Hallmark MYC targets v.1 in the Molecular Signature Database (Liberzon et al., 2015), which is compiled from previous studies. Therefore, samples with the highest activity score had, on average, the highest expression of MYC target genes. MYC activity was significantly higher in breast tumors compared to adjacent normal tissues, suggesting our MYC activity score is an appropriate method of classifying samples (Figure S1B).

We then classified MYC activity across tumor subtypes. Basal breast tumors, of which 77% are classified as triple negative (Alluri and Newman, 2014), had the highest MYC activity compared to other subtypes (Figure S1C). High levels of MYC protein and MYC-driven pathways in basal tumors versus other subtypes have been observed in other cohorts (Chandriani et al., 2009; Fallah et al., 2017), further validating our classifier. We defined MYC-active and -inactive tumors as those having a MYC-activity z-score > 1.5 and < -1.5, respectively, resulting in 78 MYC-active and 74 MYC-inactive breast tumors out of 1,073 (Figure 1A, S1D and Table S1B).

**Figure 1.**
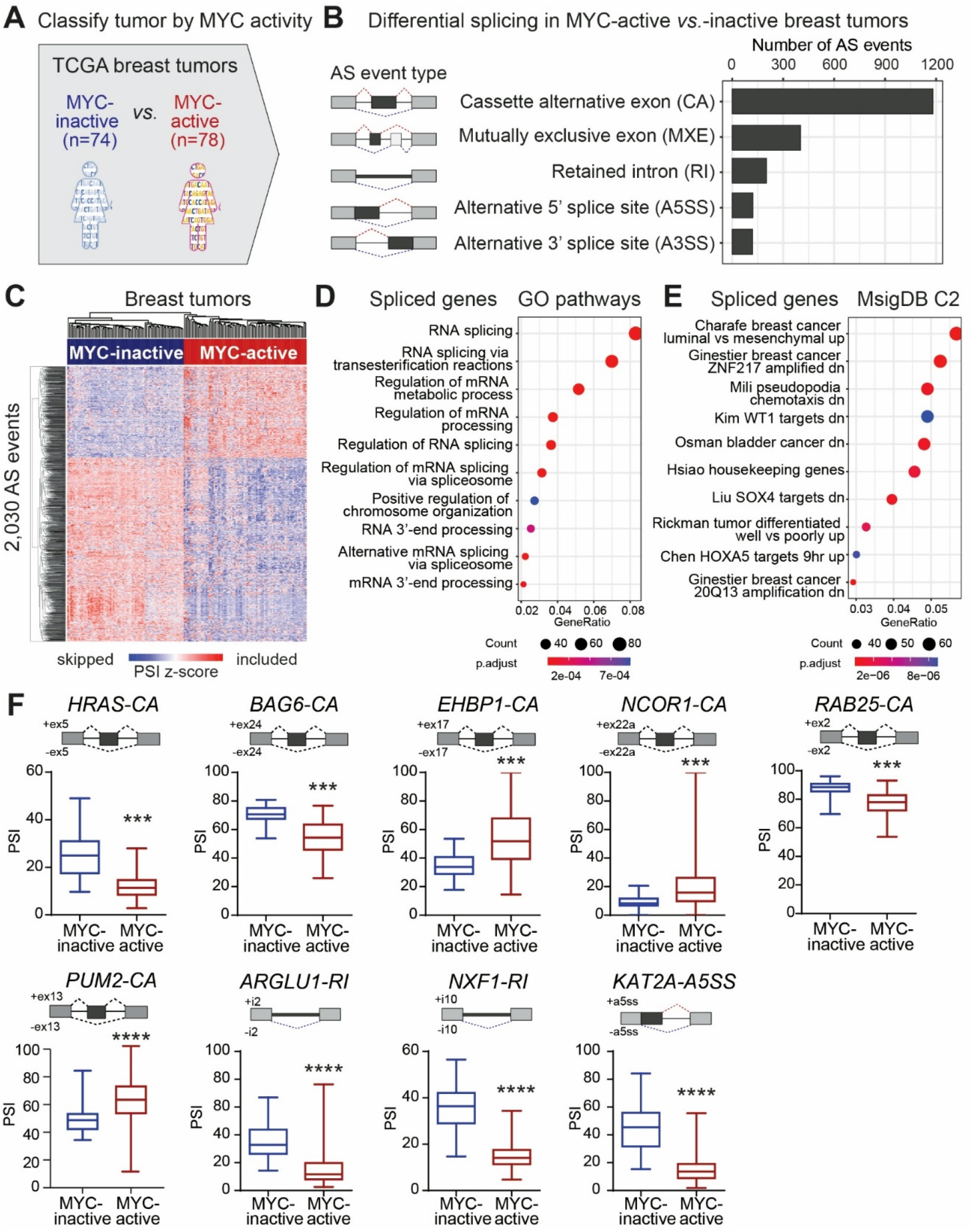
MYC-active breast tumors exhibit a unique splicing signature. **(A)** TCGA breast tumors (n=1,073) are classified by MYC activity, calculated based on the expression of known MYC targets from GSEA MSigDB Hallmark MYC targets V1 gene list (see Methods). MYC-active tumors (n=78) are defined as z-score>1.5 and MYC-inactive (n=74) tumors as z-score<-1.5. **(B)** Significant AS events detected in MYC-active *vs.* -inactive breast tumors (ΔPSI>|10%|, FDR<0.05), shown per AS event type. **(C)** Hierarchical clustering of 2,030 AS events in MYC-active (n=78) and -inactive (n=74) breast tumors. Each column represents one tumor and each row represents the percent-spliced-in (PSI) value of a specific AS event, normalized as a Z-score PSI across all tumors. **(D,E)** Gene ontology analysis using GO biological pathways gene sets (D) and MSigDB C2 chemical and genetic perturbations signature (E) for differentially spliced genes between MYC-active and -inactive breast tumors. **(F)** Percent spliced in (PSI) values of differential AS events detected in MYC-active (n=78) vs. -inactive (n=74) breast tumors (median±interquartile range; t-test,\*\*\**P*<0.0001). Gene name and AS event types are indicated. See also Figure S1 and Table S1.

We characterized the alternative splicing (AS) profiles in MYC-active *vs.* -inactive TCGA breast tumors using an in-house computational pipeline that leverages STAR for transcript assembly (Dobin et al., 2013), StringTie for reference-guided transcriptome reconstruction to identify novel spliced isoforms (Pertea et al., 2015), and rMATS for quantification of differential splicing events (Shen et al., 2014) (Figure S1E). This pipeline was built using Nextflow, a framework for data-intensive pipelines which allows for complex parallel workflows on the cloud and/or high-performance computer clusters, and utilizes containers that make installation easy and results highly reproducible. To enable efficient and cost-effective analysis of hundreds of samples simultaneously, we ran the pipeline in the cloud using Lifebit’s CloudOS platform with GCloud (see Methods). Splicing quantification was performed using rMATS at the event level using both exon body and exon junction reads. For each AS event, a percent spliced-in (PSI) value, measuring reads supporting exon inclusion *vs*. all the reads covering that event, was derived (Shen et al., 2014). We identified 2,030 differential AS events in >1,000 genes between MYC-active and -inactive breast tumors that exhibit at least 10% of change in PSI at an FDR<0.05 cutoff (Figures 1B and Table S1C). Cassette alternative exons were the most common AS event type, followed by mutually exclusive exons, retained introns, and alternative 5’ or 3’ splice sites. Principal component analysis (PCA) of the top variable AS events showed clustering of tumors based on MYC activity (Figure S1F). These 2,030 AS events provide a splicing signature that distinctly defines MYC-active *vs.* -inactive tumors (Figure 1C). Gene ontology analysis of the genes in the signature revealed an enrichment for genes associated with RNA splicing and processing, breast cancer, epithelial-to-mesenchymal transition, and chemotaxis (Figures 1D-E).

MYC-active tumors exhibit splicing changes in known cancer genes, such as skipping of a cassette exon in the *HRAS* oncogene (Figure 1F). This exon skipping event leads to the production of the longer HRAS p21 tumorigenic isoform instead of the truncated HRAS p19 isoform which lacks the ability to bind GTP and could act as a tumor suppressor (Camats et al., 2009; Cohen et al., 1989; Guil et al., 2003). This shift in expression from the p19 to the p21 isoform may contribute to the oncogenicity of MYC, and has recently been associated with MYC expression in prostate cancer (Phillips et al., 2020). Another differential AS event was detected in the *BAG6* gene, which is involved in apoptosis and ubiquitin-mediated metabolism and degradation (Kuwabara et al., 2015). MYC-active tumors display increased skipping of exon 24 (Figure 1F) which codes for the majority of the BAG-similar domain, a key mediator in the interaction of BAG6 with Ubl4A. This interaction is crucial for promoting tail-anchored transmembrane protein biogenesis and maintaining newly synthesized proteins in an unfolded state (Kuwabara et al., 2015). Therefore, skipping of exon 24 may disrupt normal functioning of BAG6, preventing it from protecting cells from misfolded protein accumulation. Other AS genes associated with MYC activity are involved in cellular organization (*EHBP1*), transcriptional regulation and/or chromatin remodeling (*NCOR1, KAT2A*), membrane trafficking and cell survival (*RAB25*), and RNA processing (*PUM2, ARGLU1, NXF1*) (Figure 1F). While some AS events are predicted to disrupt exons encoding known protein domains, including those in *HRAS*, *BAG6*, *NCOR1, PUM2*, and *KAT2A*, leading to protein isoforms with potentially distinct biological functions, other events, such as in *RAB25, ARGLU1*, or *NXF1*, introduce a premature termination codon and are predicted to decrease total protein levels. Some AS events, such as in *EHBP1*, do not disrupt the reading frame or any known domain, and therefore their functional roles are more challenging to predict.

In sum, MYC activation in breast tumors results in the expression of a distinct set of alternative isoforms that could play a role in MYC-induced oncogenesis, and includes events predicted to lead to misregulation of known cancer genes and of the RNA splicing pathway.

### Splicing factor co-expression modules correlate with MYC activity in breast tumors

To identify the SFs regulating the AS events identified above, we performed differential gene expression analysis in MYC-active *vs.* -inactive TCGA breast tumors and specifically looked for changes in SF levels. Differential gene expression analysis identified 140 upregulated and 23 downregulated SFs in MYC-active *vs*. -inactive breast tumors (Figure 2A and Table S2A). This represents a large fraction of the >300 SFs and other RNA binding proteins known to regulate splicing as defined by the ENCODE RNA consortium (Consortium, 2012; Consortium et al., 2020; Van Nostrand et al., 2020a; Van Nostrand et al., 2020b), further demonstrating MYC’s extensive role as a regulator of splicing in cancer.

**Figure 2.**
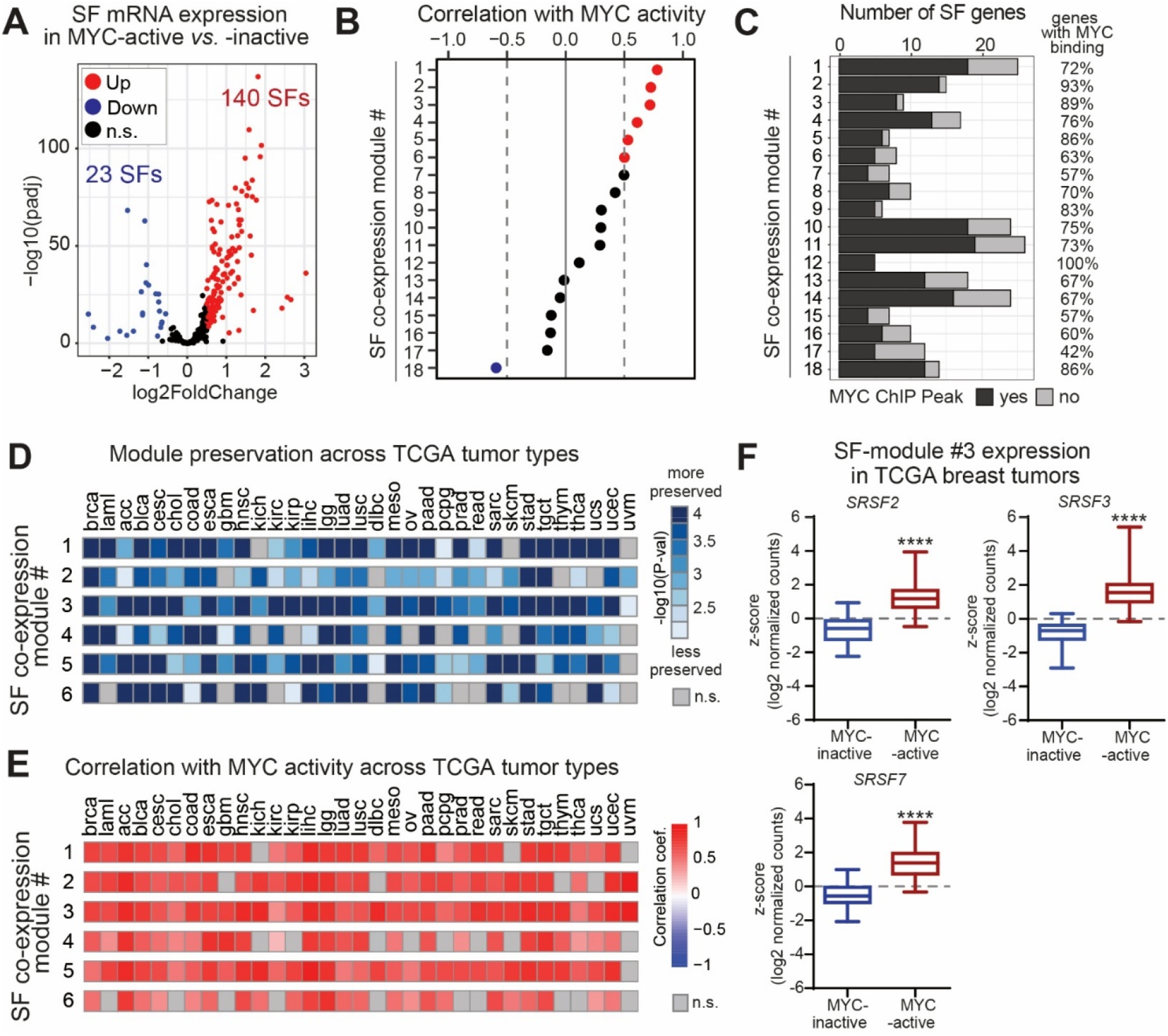
SF co-expression modules correlate with MYC activity in breast tumors and across multiple cancer types. **(A)** Differential expression of SFs in MYC-active (n=78) *vs.* -inactive (n=74) TCGA breast tumors (log_2_FoldChange>|0.5|; FDR<0.05). **(B)** Correlation of SF-module expression with MYC activity in TCGA breast tumors, shown for each module. SF-modules with correlation coefficient r>0.5 and r<-0.5 are shown in red and blue respectively. **(C)** Total number of genes in each SF-module and number of genes that exhibits at least one MYC ChIP-seq binding peak within the promoter region (<1kb from start site) in human mammary MCF-10A cells (ENCODE ENCSR000DOS) (Davis et al. 2018; Consortium 2012; Lou et al. 2020; Kim et al. 2015). The percent of genes in each module that exhibits a MYC ChIP-seq binding peak is indicated on the right. **(D)** Preservation of the top six SF-modules identified in breast tumors across 33 TCGA tumor types (see tumor names in Table S2E), represented as -log10(P-value) for the average correlation statistic using NetRep (Ritchie et al. 2016). This statistic measures the average magnitude of the correlation coefficients, or in other words, how correlated the module is on average. **(E)** Correlation coefficient of the top six SF-modules identified in breast tumors with MYC activity across 33 TCGA tumor types. **(F)** Expression of pan-cancer SF-module hub genes *SRSF2*, *SRSF3*, and *SRSF7* in MYC-active (n=78) *vs.* -inactive (n=74) TCGA breast tumors (median±interquartile range; t-test, \*\*\*\**P*<0.0001). See also Figure S2 and Table S2.

To address one of our central questions of whether SFs may function as a coordinated network regulated by MYC, we performed weighted gene correlation network analysis (WGCNA) that uses hierarchical clustering and co-expression networks to identify modules of co-expressed genes (Langfelder and Horvath, 2008, 2012). WGCNA on the ∼300 known SF genes identified 18 SF-modules, ranging in size from 5 to 26 genes, in 1,073 TCGA breast tumors (Figures 2B-C and Table S2B). For each SF-module, the most representative genes, which we call “hub” genes, were selected based on module membership correlation (MM>0.75), a quantification describing how similar the expression level of a given gene is compared to that of its module across the 1,073 tumors (Table S2B). All modules except two (modules #17 and #18) were more highly expressed in MYC-active *vs.* -inactive breast tumors (Figure S2A). We next evaluated the correlation between SF-module expression and MYC activity. The expression of SF-modules #1 through #6 was significantly positively correlated with MYC activity (r>0.5), while module #18 was negatively correlated (r<-0.5) (Figure 2B). The top three SF-modules displayed stronger correlation with MYC activity (r>0.7) than any of four individual SF genes known to be bound and directly regulated by MYC, *i.e.*, *TRA2β* (r=0.55), *SRSF1* (r=0.52), *HNRNPA1* (r=0.3), or *PRMT5* (r=0.3) (Tables S2C-D) (Das et al., 2012; David et al., 2010; Koh et al., 2015; Park et al., 2019).

To assess whether SF-module genes are direct targets of MYC, we mapped MYC chromatin immunoprecipitation sequencing (ChIP-seq) binding peaks in their promoter regions using publicly available ENCODE data from human mammary MCF-10A cells (Consortium, 2012; Davis et al., 2018; Lou et al., 2020). Over 50% of genes in each module contained at least one MYC binding peak in their promoter region, with the exception of module #17 with 41% (Figure 2C and Table S2B). Modules #2, #3, #5, #12, and #18 had >85% of genes with MYC binding peaks in their promoter regions, suggesting they are likely directly transcriptionally regulated by MYC.

To validate that the relationship between SF-modules and MYC activity is not limited to the TCGA cohort, we assessed the preservation and correlation of SF-module expression with MYC activity in 2,969 breast tumors from the Sweden Cancerome Analysis Network Breast (SCAN-B) cohort (Brueffer, 2018). Similar to the TCGA cohort, tumors of a basal-like subtype had significantly higher MYC activity scores than other less aggressive subtypes (Figure S2B). To determine whether SF-modules detected in the TCGA cohort were preserved in the SCAN-B cohort, we analyzed all tumors in the SCAN-B cohort and utilized NetRep, a computational tool that performs permutation tests to evaluate the preservation of co-expression modules across datasets, focusing on four parameters recommended for small module sizes (see Methods) (Ritchie et al., 2016). A module was classified as preserved if three out of four preservation statistics reached a threshold p-value ≤ 0.01. All but one of the 18 SF-modules were preserved across SCAN-B tumors (Figure S2C-D). Of these, 10 SF-modules had a strong correlation with MYC activity in the SCAN-B cohort (Figure S2E). In particular, SF-modules #1-6 were preserved and correlated with MYC activity across both cohorts (Figures 2D and S2E), further supporting that MYC regulates their expression in breast cancer.

In sum, across two independent breast cancer cohorts, MYC activity correlates with the expression of SF-modules, the majority of which are directly bound by MYC in their promoter regions. Therefore, the expression of these SF-modules in breast tumors may contribute to the difference in splicing patterns observed between MYC-active and -inactive tumors (Figure 1).

### SF-module pan-cancer preservation and correlation with MYC activity

We next assessed whether the MYC-regulated SF-modules identified in breast tumors were preserved in other tumor types, focusing on six SF-modules which had a correlation with MYC activity greater than 0.5 in both breast cancer cohorts. We scored MYC activity across 32 additional TCGA tumor types and used NetRep to determine the preservation of each module as above. We found one module - SF-module #3 – that was preserved in all 32 tumor types, and even the least preserved SF-module #6 was found in 23 other cancers (Figures 2D and S2F). All six SF-modules maintained a strong correlation (r>0.5) with MYC activity in those tumor types in which they were significantly preserved (Figure 2E), suggesting that MYC regulates expression of these modules across tumor types. The hub genes of the pan-cancer SF-module #3, *SRSF2*, *SRSF3*, and *SRSF7,* are all members of the SR protein family, a group of SFs which have been previously found dysregulated in various tumor types (Fu and Wang, 2018; Jia et al., 2010; Luo et al., 2017; Park et al., 2019; Saijo et al., 2016; Song et al., 2019; Urbanski et al., 2018; Wang et al., 2020a). These three SR protein genes are each significantly upregulated (∼1.5 to 2-fold) in MYC-active tumors compared to MYC-inactive tumors in both TCGA and SCAN-B breast cohorts (Figures 2F and S2G), and contain ChIP-seq MYC binding peaks (Table S2B).

### MYC activation induces alternative splicing changes and upregulates the pan-cancer SF-module in human mammary epithelial cells

We aimed to further experimentally validate that the SF-modules preserved across cancers and AS events detected in TCGA tumors are regulated by MYC. To study the effects of MYC activation on SF-module expression and splicing in a controlled inducible model system, we used non-transformed human mammary epithelial MCF-10A cells expressing MYC fused to a portion of estrogen receptor (MYC-ER) (Eilers et al., 1989; Littlewood et al., 1995). MYC is activated by treating the cells with 4-hydroxytamoxifen (4-OHT), which promotes translocation of the MYC-ER fusion protein into the nucleus and induces transcription of MYC target genes. To determine the rate of MYC induction after 4-OHT treatment and an optimal timepoint for our analysis, we assessed the mRNA and protein levels of two known MYC target genes, *SRSF1* and *TRA2β* over 48 hours (Figure S3A-B) (Anczukow et al., 2012; Park et al., 2019). Based on these results, we performed RNA-seq, in triplicate, on 3D-grown MCF-10A MYC-ER cells at 0, 8, and 24 hours (h) after 4-OHT-induced MYC activation. As a control for 4-OHT-induced effects, 3D-grown parental MCF-10A cells lacking the MYC-ER fusion protein were treated with 4-OHT at the same timepoints. PCA using gene expression counts showed robust clustering of the MCF-10A MYC-ER samples by MYC-activation timepoint, whereas in MCF-10A controls, 4-OHT had minor effects on gene expression (Figure S3C). MYC activation was confirmed by assessing the expression of our panel of 200 known MYC target genes (Liberzon et al., 2015) (Figure S3D).

We first addressed whether MYC activation in MCF-10A cells induced expression of SFs. We identified 138 and 119 upregulated SFs at 8h and 24h of MYC activation, respectively, and 28 and 21 downregulated SFs at 8h and 24h, respectively (Figure 3A-B and Table S3A,B). 121 SFs were differentially expressed and regulated in the same direction between 8h and 24h (Figure S3E). Comparison with TCGA breast tumors revealed 90 SFs differentially expressed in all three MYC-active *vs.* -inactive conditions, and only 3 SFs changed in opposing directions between TCGA breast tumors and MCF-10A MYC-ER cells (Figure 3C). These data suggest that these 90 SFs are MYC-regulated in breast cells and tumors.

**Figure 3.**
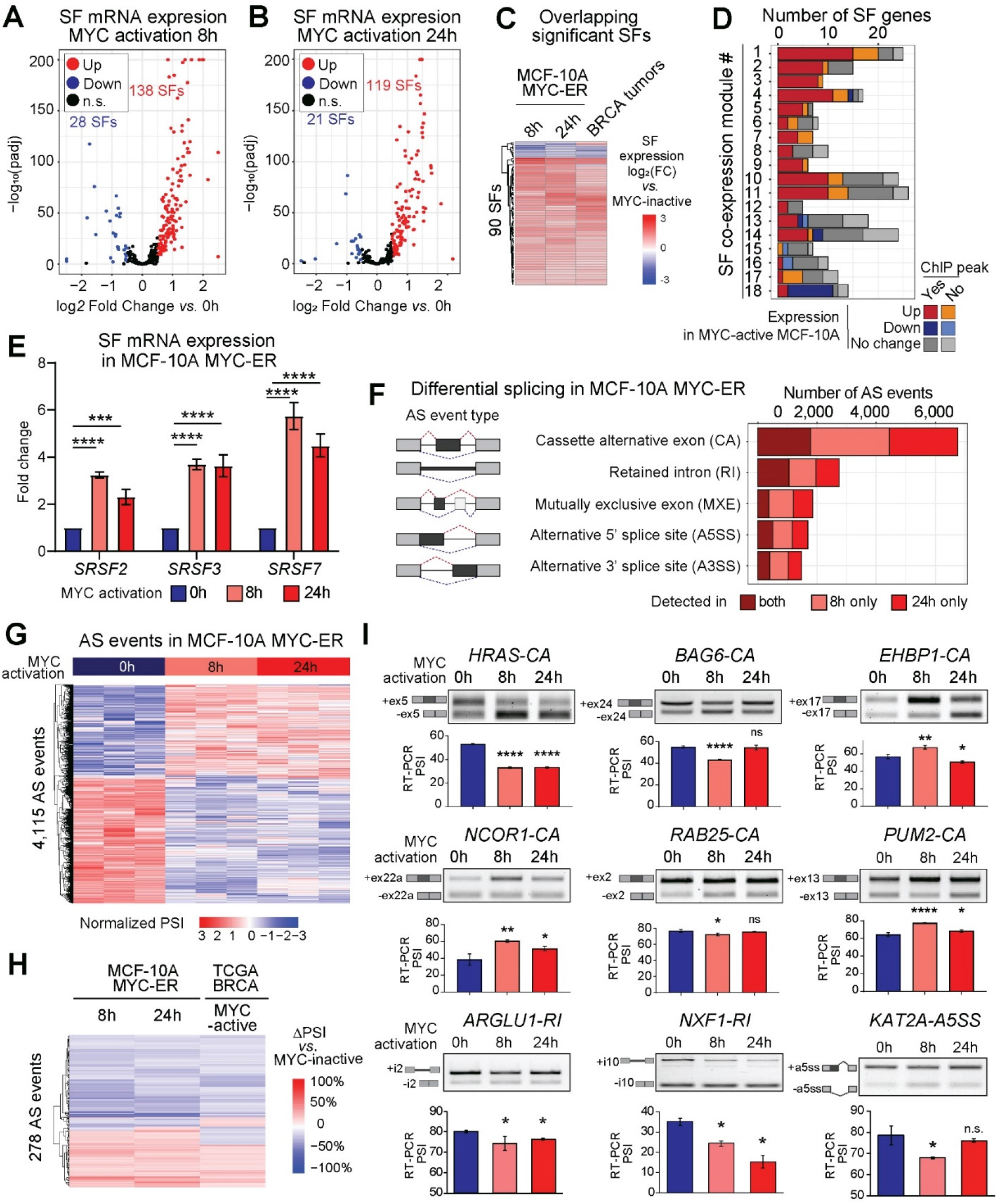
MYC activation induces changes in SF expression and alternative splicing in human mammary epithelial cells. **(A,B)** Differential expression of SFs in MCF-10A MYC-ER cells at 8h (A) or 24h (B) *vs.* 0h after MYC activation (n=3; log_2_FoldChange>|0.5|; FDR<0.05). The number of significantly up-regulated, down-regulated genes in MYC-active *vs.* -inactive is indicated (n.s. – not significant). **(C)** Differential expression of 90 SFs shared across MYC-active MYC-ER cells at 8h, 24h, and TCGA breast tumors, as compared to MYC-inactive cells and tumors (log_2_FoldChange>|0.5|; FDR<0.05). **(D)** Number of genes for each TCGA-derived SF-module differentially expressed in MCF-10A MYC-ER after MYC activation, and number of genes per SF-module that exhibits at least one MYC ChIP-seq binding peak within the promoter region (<1kb from start site) (Davis et al. 2018; Consortium 2012; Lou et al. 2020; Kim et al. 2015). **(E)** mRNA expression of pan-cancer SF-module hub genes *SRSF2*, *SRSF3*, and *SRSF7* in MCF-10A MYC-ER cells after MYC activation, normalized to 0h, measured by Q-PCR using gene specific primers, normalized to GAPDH (n=3; mean±sd; two-way ANOVA;\*\**P*<0.005, \*\*\*\**P*<0.0001). **(F)** Number of significant AS events detected in MCF-10A MYC-ER cells at 8h or 24h *vs.* 0h after MYC activation (n=3 per condition; |ΔPSI|≥10%; FDR<0.05) show per event type. **(G)** Hierarchical clustering of significant AS events detected in MCF-10A MYC-ER cells at both 8h and 24h *vs.* 0h after MYC activation (n=3 per condition; |ΔPSI|≥10%; FDR<0.05). Each column represents one AS event and each row represents the percent-spliced-in (PSI) value of a specific AS event, normalized as a Z-score PSI across all samples.**(H)** Difference in percent-spliced-in (ΔPSI) for 278 significant AS events detected in MCF-10A MYC-ER cells 8h or 24h *vs.* 0h, and MYC-active *vs.* inactive TCGA breast tumors (|ΔPSI|≥10%; FDR<0.05). **(I)** Validation of selected MYC-regulated AS events in MCF-10A MYC-ER cells at 8h or 24h *vs.* 0h after MYC activation by RT-PCR using primers that amplify included and skipped events. A representative gel is shown, along with isoform structures. Splicing quantification as PSI from RT-PCR are shown (n=3; mean±sd; t-test, \**P*<0.05, \*\**P*<0.01, \*\*\**P*<0.001, \*\*\*\**P*<0.0001, n.s. – not significant). Gene name and AS event types are indicated. See also Figure S3 and Table S3.

We next assessed whether the SF co-expression modules identified in TCGA breast tumors and preserved in others were also differentially expressed in MCF-10A MYC-ER cells after MYC activation. In the SF-modules most associated with MYC activity in the TCGA cohort (Figure 2B), such as SF modules #1-6, at least 50% of the genes were upregulated by MYC after either 8h or 24h (Figure 3D), and the majority also exhibited MYC Chip-seq binding peaks within their promoter regions. Modules which did not strongly correlate with MYC activity in TCGA and SCAN-B tumors, such as modules #12-17 (Figure 2B and S2E), had fewer than 50% of SF genes upregulated in MCF-10A MYC-ER cells. Module #18, which was significantly anti-correlated with MYC activity in TCGA breast tumors, had 65% genes downregulated. Thus, we see a remarkable overlap between our *in vitro* model and TCGA breast tumors in terms of SFs associated with MYC activity. Given the pan-cancer preservation and correlation of SF-module #3 with MYC activity, we aimed to further investigate whether these genes are directly induced by MYC and whether they regulate splicing in a cooperative manner. Pan-cancer SF-module #3 hub genes, *SRSF2*, *SRSF3*, and *SRSF7,* exhibit experimental evidence of MYC binding in their promoter regions in ChIP-seq experiments in MCF-10A cells (Figure S3F and Table S2B), and expression of all three genes increases after MYC activation as measured by quantitative PCR (qPCR) (Figure 3E).

We then performed differential splicing analysis, as described above, comparing MYC-active (8h or 24h) *vs*. inactive (0h) MCF-10A MYC-ER cells. MCF-10A cells lacking the MYC-ER fusion protein were used to control for 4-OHT-induced effects; a small number of AS events were detected in 4-OHT treated-MCF-10A control cells, and were removed from further analysis. We identified >9,000 significant AS events in >4,000 genes after 8h of MYC activation and >8,000 significant AS events in >3,000 genes after 24h of MYC activation compared to 0h, with at least a 10% change in PSI at an FDR<0.05 cutoff (Figure 3F and Tables S3C,D). Approximately 40% of the AS events detected at 8h were also detected at 24h, the majority of which changed in the same direction (*i.e*., included or excluded) (Figures 3G, S3G and Tables S3E,H). Most MYC-induced AS events were cassette alternative exon events, the most common AS events, followed by retained introns and mutually exclusive exons. Genes alternatively spliced after 8 or 24h of MYC activation were most enriched in RNA processing and metabolism processes as well as cancer-related pathways (Figure S3H-K).

Finally, we compared the AS events detected in MYC-active MCF-10A MYC-ER mammary cells with those detected in MYC-active breast tumors. We found 706 AS events that were differentially spliced in both TCGA MYC-active tumors and MCF-10A MYC-ER activated for either 8h or 24h (Tables S3F-H). There were 278 AS events detected in all three datasets, of which >80% changed in the same direction (Figure 3H). We detected more MYC-induced AS events in MYC-active mammary cells compared to tumors, likely due to the higher RNA-sequencing depth in cell lines compared to tumors (on average >100M vs. >30M reads per sample), as well as to the genomic and transcriptomic heterogeneity of tumors which renders the detection of recurring AS events across multiple whole tumor samples more difficult. We validated 17 AS events in MCF-10A MYC-ER cells via RT-PCR (Figures 3I and S3L) involved in cancer-related processes such as cellular organization (*EHBP1*), transcriptional regulation and/or chromatin remodeling (*NCOR1, KAT2A*), membrane trafficking and cell survival (*RAB25*), RNA processing (*PUM2, ARGLU1, NXF1*), signal transduction (*APLP2, FAM126A, HPS1*), metabolism (*BTN2A1*, *LSR*), cell cycle (*CENPX*), autophagy (*WDR45*), or vesicle trafficking (*TEPSIN*). Of these, 9 were also detected in TCGA MYC-active breast tumors (Figure 1F), including skipping of the cassette alternative exon 5 in the *HRAS* oncogene.

### Pan-cancer SF-module hub genes, SRSF2, SRSF3, and SRSF7, control splicing of MYC-regulated target exons

We next sought to determine which MYC-regulated AS events detected in both MCF-10A MYC-ER cells and TCGA breast tumor samples are likely to be directly regulated by the hub genes SRSF2, SRSF3 and SRSF7 of the pan-cancer SF-module #3. SR proteins regulate splicing by binding specific motifs on the pre-mRNA and promoting exon inclusion or skipping (Long and Caceres, 2009). Although SR proteins have evolved from a common ancestor, exhibit motif similarities, and share some targets, they can also regulate distinct sets of splicing events (Bradley et al., 2015; Long and Caceres, 2009; Park et al., 2019; Van Nostrand et al., 2020a). We first mapped known SRSF2, SRSF3, and SRSF7 binding motifs (Figures 4A-C and S4A) in 6 of the validated MYC-regulated AS events (Figure 3I). SRSF2, SRSF3 and SRSF7 binding motifs were derived from *in vitro* and *in vivo* experiments including Systematic Evolution of Ligands by Exponential Enrichment (SELEX), RNAcompete, or RNA Bind-n-Seq (Table S6A) (Ajiro et al., 2016; Dominguez et al., 2018; Kim et al., 2015; Konigs et al., 2020; Paz et al., 2014; Ray et al., 2013), and mapped using RBPmap (Paz et al., 2014). Motif analysis revealed that all 6 analyzed events exhibit motifs for all 3 of SRSF2, SRSF3 and SRSF7 either in the alternative exon or surrounding introns. We found evidence for overlapping motifs as well as for non-overlapping motifs for each event, both in exonic and intronic regions (Figures 4A-C and S4A). These findings suggest that SRSF2, SRSF3 and SRSF7 likely bind and regulate the splicing of these exons, either by binding at the same time, by competing for binding at overlapping motifs, or a combination of these two mechanisms.

**Figure 4.**
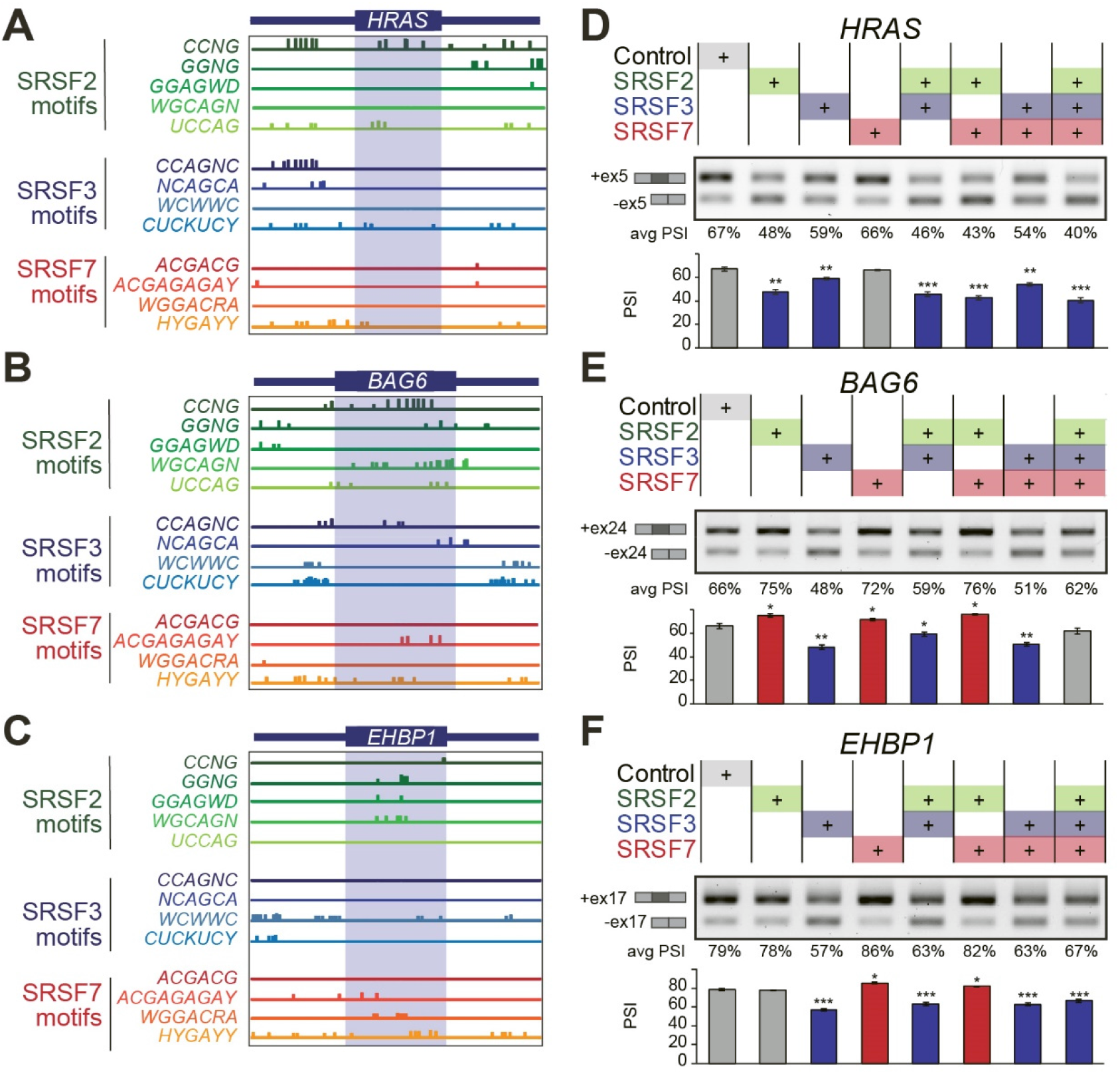
MYC-regulated splicing events display binding motifs for and are regulated by pan-cancer SF-module hub genes SRSF2, SRSF3, and SRSF7. **(A-C)** Position of predicted binding motifs for SRSF2, SRSF3, or SRSF7 in the spliced sequences (blue box) and surrounding introns (100bp) of validated MYC-regulated AS events in MCF-10A cells. Consensus motif sequences derived from *in vitro* or *in vivo* studies are indicated (Ajiro et al., 2016; Dominguez et al., 2018; Kim et al., 2015; Konigs et al., 2020; Paz et al., 2014; Ray et al., 2013). **(D-F)** Splicing of MYC-regulated splicing events in HEK293 cells transfected with the coding sequence for either one, two, or all three SR proteins, or control plasmids, is measured by RT-PCR using primers that both amplify included and skipped isoforms. A representative gel is shown, along with isoform structures. Splicing quantification as PSI from RT-PCR (n=3; mean±sd; t-test, \**P*<0.05, \*\**P*<0.001, \*\*\**P*<0.0001). See also Figures S4-S5 and Table S6A

We next sought mechanistic insight into how SRSF2, SRSF3 and SRSF7 impact splicing of MYC-regulated events and whether these SFs exhibit cooperative or antagonistic effects. We co-transfected one, two, or all three HA-tagged plasmids encoding for these SR proteins, or a control plasmid, into HEK293T cells and measured splicing of 6 breast cancer-relevant MYC-regulated events from Figure 3I by RT-PCR (Figures 4D-F and S4B,C). We uncovered some events that are regulated in the same direction by multiple SR proteins, such as the *HRAS* event. Indeed, expression of SRSF2 alone decreased *HRAS* exon 5 inclusion (PSI 47%), while SRSF3 had a milder effect (PSI 59%), and SRSF7 had no effect (PSI 66%) compared to control (PSI 67%) (Figure 4D). Co-expression of SRSF2 together with SRFS7 promoted even more skipping (PSI 43%) compared to either SF alone, while SRSF2 and SRSF3 together (PSI 46%) had no stronger effect than either one alone. Co-expression of all three SFs led to the strongest exon 5 skipping effect in *HRAS* (PSI 40%). Conversely, SR proteins can also regulate splicing events in opposite directions, such as those in *BAG6* and *EHBP1* (Figure 4E-F). While SRSF3 expression decreased *BAG6* exon 24 inclusion (PSI 48%), expression of SRSF2 or SRSF7 increased exon inclusion (PSI 75% and 72%) compared to control (PSI 66%) (Figure 4E). Co-expression of SRSF3 and SRSF7 decreased exon 24 skipping (PSI 59%) compared to SRSF3 alone. In the case of *EHBP1,* exon 17 inclusion was positively regulated by SRSF7 (PSI 86%) and negatively by SRSF3 (PSI 57%), while SRSF2 had no effect (PSI 78%) compared to control (PSI 79%) (Figure 4F). Co-expression of SRSF3 and SRFS7 decreased exon 17 inclusion (PSI 63%) compared to SRSF3 alone (Figure 4F). These results demonstrate that the action of SR proteins in regulation of alternative exons can be cooperative, such as SRSF2 and SRSF3 in *HRAS*, or competitive, such as SRSF3 and SRSF7 in *BAG6* and *EHBP1*.

We also aimed to determine whether these MYC-regulated AS events are controlled by MYC-regulated pan-cancer SF-module #3 in established breast cancer cells. Using publicly available RNA-seq data from the Cancer Cell Line Encyclopedia (CCLE) (Ghandi et al., 2019), we ranked all breast cancer cell lines by MYC activity score and selected MDA-MB231 cells as a representative MYC-active triple negative basal breast cancer cell line (Figure S5A). We generated stable cell lines expressing a tetracycline-regulated transactivator (rTTA3), along with doxycycline (DOX)-inducible shRNAs targeting each of the three hub genes from pan-cancer SF-module #3, SRSF2, SRSF3, SRFS7, or control (Figure S5B). We evaluated the effect of knockdown of each SR protein individually in MDA-MB231 cells on splicing of known MYC targets previously identified in breast tumors. Knockdown of SRSF3, but not SRSF2 or SRSF7, increased exon inclusion in *BAG6* (Figure S5C), thus mimicking the effects of low MYC activity as detected both in TCGA tumors (Figure 1F) and MCF-10A cells (Figure 3I) and promoting the reverse splicing pattern observed in cells overexpressing SRSF3 (Figure 4E). Similarly, reduced exon inclusion of *EHBP1,* as detected in low MYC-activity cells and tumors (Figures 1F and 4E), was triggered by knockdown of SRSF7, while SRSF3 knockdown increased exon inclusion of *EHBP1* (Figure S5C). SRSF2 or SRSF3 knockdown decreased *NCOR1* exon inclusion, while SRSF7 knockdown promoted exon inclusion (Figure S5C). Finally, SRSF2 or SRSF7 knockdown promoted *PUM2* exon inclusion, while SRSF3 knockdown promoted skipping (Figure S5C). None of the knockdowns significantly impacted *HRAS* splicing (Figure S5C), which at baseline is mostly skipped in these established cancer cells.

Taken together, our data from SF-overexpression and -knockdown experiments in multiple cell models (Figures 4 and S5), suggest that these MYC-associated splicing events are regulated by pan cancer SF-module hub genes, SRSF2, SRSF3, and SRSF7. The effect of pan cancer SF-module on targets is dependent on the expression level of individual SR proteins as well as their coordinated ability to regulate splicing.

### Pan-cancer SF-module hub genes are necessary and sufficient for breast cancer organoid size and invasiveness

We next evaluated how knockdown of pan cancer SF-module hub genes impacts growth and invasion of a MYC-active breast cancer cell line that exhibits invasive and metastatic properties. In 3D-grown MDA-MB231 cells, knockdown of either SRSF3 or SRSF7 significantly decreased both the size of organoids and the presence of invasive cellular projections from each organoid (Figure 5A), with the shRNA targeting SRSF3 exhibiting the strongest phenotype. Additionally, both SRSF3 and SRSF7 knockdowns slightly decreased cell proliferation, while SRSF2 had no effect (Figure S5D). SRSF2 knockdown had minimal effects on measured cellular phenotypes, although this could be due, at least in part, to a milder knockdown efficiency with the SRSF2-targeting shRNA (Figure S5B). Overall, knockdown of either SRSF3 or SRSF7 mimicked the effect of MYC knockdown in this model (Figure 5A and S5D), suggesting that these two hub genes are required for invasion in this MYC-active cell line.

**Figure 5.**
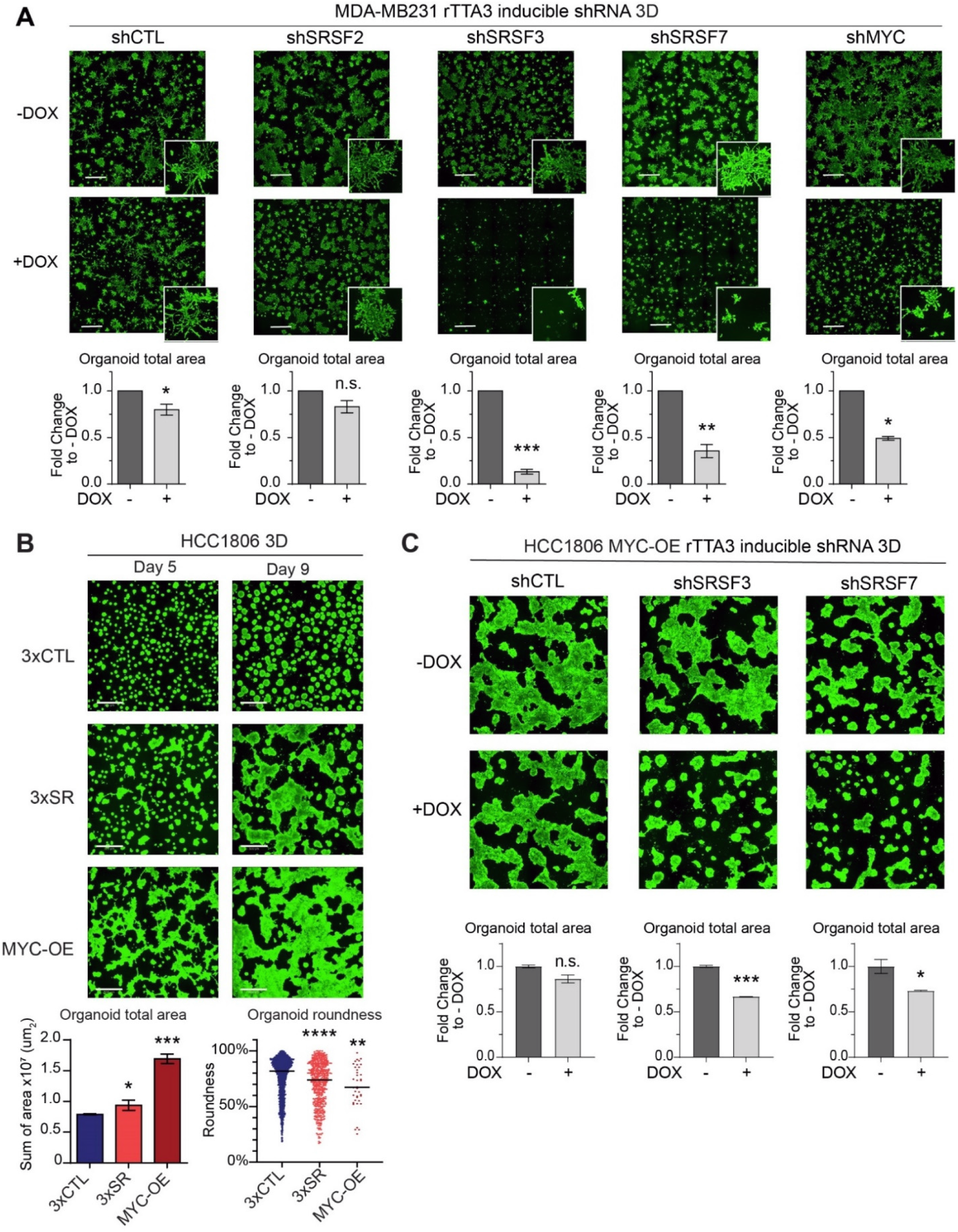
Pan-cancer SF-module hub genes SRSF2, SRSF3, and SRSF7 control breast cancer organoid size and invasiveness. **(A)** Representative images of 3D-grown MDA-MB231-rTTA3 cells expressing DOX-inducible shRNA targeting SRSF2, SRSF3, SRSF7, MYC, or control (CTL) at day 9 stained with calcein (scale bar: 1mm). For each condition, >30 fluorescent confocal z-stacks images are combined as a maximal projection image. Total organoid area is quantified and plotted compared to -DOX control for each condition (n=3, 25 fields per replicate; mean±sd; t-test, \**P*<0.05, \*\**P*<0.01, \*\*\**P*<0.0001, n.s.-not significant). **(B)** Representative images of 3D-grown 3xCTL, 3xSR (SRSF2-SRSF3-SRSF7), and MYC-OE HCC1806 organoids at days 5 and 9 stained with calcein (scale bar: 500μm). >20 fluorescent confocal z-stacks are combined as a maximal projection image. Total organoid area (n=3, 15 fields per replicate; mean±sd; t-test, \**P*<0.05, \*\*\**P*<0.001) and organoid roundness (n=3, 15 fields per replicate; median; t-test, \*\**P*<0.01, \*\*\*\**P*<0.0001) are quantified for each condition at day 9. **(C)** Representative images of 3D-grown MYC-OE-rTTA3 HCC1806 organoids expressing DOX-inducible shRNA targeting SRSF3, SRSF7, or control (CTL) at day 9 stained with calcein (scale bar: 500μm). For each condition, >30 fluorescent confocal z-stacks images are combined as a maximal projection image. Total organoid area is quantified and plotted compared to -DOX control for each condition (n=2, 25 fields per replicate; mean±sd; t-test, \**P*<0.05, \*\*\**P*<0.0001, n.s.-not significant). See also Figures S5-S6.

In parallel, to determine whether MYC-regulated expression of pan cancer SF-module hub genes, *SRSF2*, *SRSF3*, and *SRSF7* contributes to oncogenesis, we assayed the functional consequences of overexpressing all three genes together in MYC-inactive breast cancer cells. We selected HCC1806 as a representative MYC-inactive basal subtype breast cancer cell line (Figure S5A), and generated stable cell lines overexpressing: i) the coding sequences for all three hub genes in pan-cancer SF-module #3 combined, *i.e.*, *SRSF2*, *SRSF3*, and *SRSF7* (hereafter called 3xSR); ii) the corresponding three empty vector controls (3xCTL); or iii) the coding sequence of MYC (MYC-OE). The 3xSR cell line exhibited at least a 1.5-fold increase in expression of each of the corresponding *SRSF* transcripts (Figure S6A), which is within the levels of *SRSF2*, *SRSF3*, and *SRSF7* overexpression in MYC-active TCGA breast tumors (Figures 2F, S2G and Table S2A) and MCF-10A MYC-ER cells (Figure 3E).

We first characterized the phenotypes of 3xSR *vs.* 3xCTL HCC1806 cells in 2D culture. Co-overexpression of SRSF2, SRSF3, and SRSF7 did not confer any proliferative advantage to HCC1806 cells in 2D (Figure S6B). In fact, 3xSR cells grew slower than 3xCTL, which could reflect cell stress from overexpressing three SR proteins. However, cell migration assays revealed 3xSR and MYC-OE were significantly more invasive than 3xCTL cells in 2D (Figure S6C). In addition, both 3xSR and MYC-OE cells exhibited changes in cell morphology compared to 3xCTL cells, including increased appearance of actin-rich filopodia (Figure S6D), consistent with the increased migratory phenotype.

We next characterized cell phenotypes in 3D growth assays in Matrigel. Both 3xSR HCC1806 and MYC-OE cells formed significantly larger and less circular organoids compared to 3xCTRL cells (Figure 5B). Since the 3xSR HCC1806 cell line was generated using successive infections of plasmids containing the coding sequence for each SR protein, we evaluated the effect of each additional SR protein on 3D growth and found that each sequential SR protein increased the invasive phenotype compared to control, suggesting SRSF2, SRSF3 and SRSF7 cooperate to increase invasion of cancer organoids (Figure S6E-F). In 3D invasion assays in collagen-Matrigel matrix, 3xSR HCC1806 cells formed more invasive structures compared to 3xCTL organoids, which rarely display any invasive behavior (Figure S6G).

Finally, to evaluate how each pan-cancer SF-module hub gene contributes to MYC-dependent phenotypes, we generated MYC-OE HCC1806 cells that stably express rTTA3 along with doxycycline (DOX)-inducible shRNAs targeting the two hub genes that impacted cellular phenotypes in MDA-MB231 cells, SRSF3 and SRFS7, plus control (Figure S6H). Knockdown of either SRSF3 or SRSF7 significantly decreased organoid size in MYC-OE HCC1806 cells (Figure 5C), but neither alone is sufficient to fully reverse the MYC-induced phenotype in this model. In sum, our findings demonstrate that the hub genes from pan-cancer SF-module are sufficient to cooperatively promote cell invasion in MYC-inactive breast cancer cell lines, and required for maintenance of invasiveness in MYC-active cells.

### Co-expression of pan-cancer SF-module hub genes leads to splicing changes in breast cancer cells

We performed RNA-seq on 3xSR, 3xCTL, and MYC-OE HCC1806 cells, in triplicate, and performed differential splicing analysis. PCA of gene expression counts showed replicates clustering based on sample (Figure S7A). At the gene expression level, in concordance with increased cell migration and invasion (Figures 5 and S6), both 3xSR and MYC-OE cells increased expression of the mesenchymal cell marker vimentin, and EMT-inducing transcription factors *TWIST1*, *TWIST2*, and *SNAI1* compared to 3xCTL HCC1806 cells (Figure S7B-D). At the splicing level, we identified >5,000 AS events in >3,000 genes in 3xSR HCC1806 cells, and >7,000 AS events in >3,000 genes in MYC-OE HCC1806 cells, compared to 3xCTL (Figure 6A-B and Tables S4A-B). Most AS events were cassette alternative exons, followed by intron retention. >1,000 AS events were differentially spliced in both 3xSR-OE and MYC-OE HCC1806 cells (*P* <2.2E-16), of which 94% changed in the same direction (Figure 6C and Table S4C,L), suggesting that SRSF2, SRSF3, and SRSF7 regulate a significant subset of AS events downstream of MYC in HCC1806 cells. Spliced genes were enriched in similar pathways in both 3xSR and MYC-OE, including processes and gene sets associated with breast cancer, chemotaxis, cellular transport and organization (Figures 6D-E and S7E-F). We validated 5 events by semi-quantitative RT-PCR (Figure S7I). One example is the skipping of exon 11a in *ENAH*, also known as Mena, which regulates actin polymerization and modulates cell motility. Expression of the exon 11a-included isoform, Mena11a, has been associated with epithelial cancer cells and decreased inclusion of exon 11a is associated with mesenchymal markers and invasion (Balsamo et al., 2016). HCC1806 3xSR and MYC-OE both had decreased inclusion of exon 11a in *ENAH*, which may contribute to increased invasiveness in these cell lines. We also validated a splicing event in the RNA binding protein *PUM2* (Figure S7I), additionally detected in MCF-10A MYC-ER activated cells (Figure 3I) and MYC-active TCGA breast tumors (Figure 1E). PUM2 has been previously implicated in promoting stemness of breast cancer cells as well as proliferation and migration in glioblastoma (Wang et al., 2019b; Zhang et al., 2019). We validated other splicing events in genes involved in hormone-mediated transcription and alternative splicing (*RBM39*), toll-like receptor signaling (*RNF216*) and cell adhesion and invasion (*PVR* also known as *CD155*) (Figure S7I).

**Figure 6.**
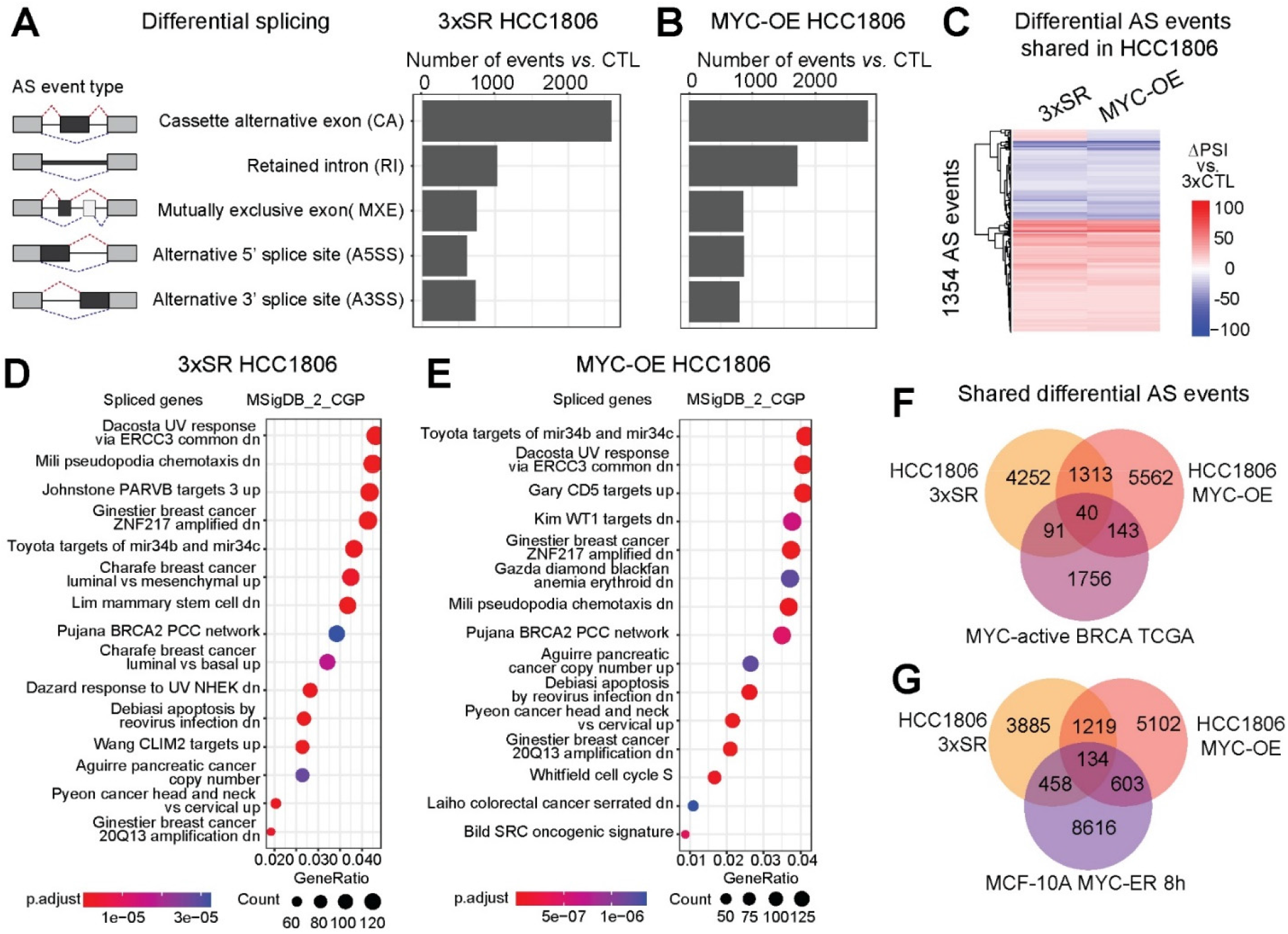
Overexpression of pan-cancer SF-module hub genes SRSF2, SRSF3, and SRSF7 together leads to splicing changes in MYC-regulated splicing events. **(A,B)** Number of significant AS events detected in 3xSR (SRSF2-SRSF3-SRSF7) (A) or MYC-OE (B) *vs.* 3xCTL HCC1806 cells (n=3 per condition; |ΔPSI|≥10%; FDR<0.05) shown per event type. **(C)** Overlapping significant AS events in 3xSR and MYC-OE *vs*. 3xCTL HCC1806 cells. Each row represents one AS event. **(D,E)** Gene ontology analysis using MSigDB C2 chemical and genetic perturbations signature for differentially spliced genes in 3xSR (D) and MYC-OE (E) *vs.* 3xCTL HCC1806 cells. **(F,G)** Overlapping significant AS events in 3xSR cells and MYC-OE HCC1806 cells and also in MYC-active TCGA breast tumors (F) or MYC-active MCF-10A MYC-ER cells (G). These are overlapping events, without considering direction of change. See also Figure S7 and Table S4.

We next investigated the extent of overlap of AS events, in addition to that in *PUM2*, between 3xSR or MYC-OE HCC1806 cells and MYC-active breast TCGA tumors. 131 differential AS events were detected in both 3xSR HCC1806 cells and MYC-active breast tumors (*P*<2.2E-16), 73% of which were changing in the same direction (Figure 6F and Table S4D,L). In comparison, 183 AS events were detected in both MYC-OE HCC1806 cells and MYC-active breast tumors (*P*=4.4E-16), of which 58% were changing in the same direction (Figure 6F and Table S4E,L). In total, there were 40 AS events that were significantly differentially spliced in all three datasets (Figures 6F, S7G and Table S4F). These findings suggest that MYC-regulated AS events detected in breast tumors and cell lines, are, at least in part, controlled by changes in SRSF2, SRSF3, and SRSF7 levels.

We then assessed the overlap of differential AS events between HCC1806 and MYC-active MCF-10A MYC-ER cells. We identified 592 AS events overlapping between 3xSR HCC1806 cells and MYC-ER 8h MCF-10A cells (*P*=2.9E-01), of which 38% were changing in the same direction (Figure 6G, Table S4G,L). In comparison, there were 737 AS events overlapping between MYC-OE HCC1806 cells and MYC-ER 8h MCF-10A cells (*P*<2.2E-16), of which 60% were changing in the same direction (Figure 6G and Table S4H,L). In total, there were 134 AS events in common between all three datasets (Figures 6G, S7H and Table S4I). Thus, a subset of MYC-induced AS events in non-transformed epithelial cells are also detected in breast cancer cells overexpressing either MYC or pan cancer SF-module hub genes, suggesting some overlapping roles for MYC in both systems. Yet, as anticipated, MYC activation in HCC1806 *vs.* MCF-10A cells also leads to differences in splicing profiles, illustrating differences in splicing regulation between non-transformed epithelial cells and established breast cancer cells.

### Pan-cancer splicing signature of MYC activity shared across multiple tumor types

Having shown the functional importance of the hub SFs in breast cancer models, we next sought to determine whether MYC-regulated AS alterations are specific to breast tumors, or if they extend to other tumor types and may also be regulated by the hub SFs therein. We thus profiled AS events correlated with MYC activity across 32 additional tumor types. As described above for breast tumors, we first divided tumors into MYC-active and -inactive groups, based on MYC activity scores, and quantified differential AS events between MYC-active *vs.* -inactive tumors for each of TCGA tumor type (Table S5). Numbers of differential AS events between MYC-active *vs.* -inactive tumors ranged from 218 for uveal melanoma (UVM) tumors to 2,549 for testicular germ cell tumors (TGCT), at |ΔPSI|≥10% and FDR<0.05 cutoffs (Figure 7A and Table S5). We first focused on splicing events associated with MYC activity in breast tumors and validated in our mammary cell models (Figures 1, 3 and 4). For example, skipping of *HRAS* exon 5, which favors the expression of tumorigenic HRAS, was significant (FDR<0.05) across 25 tumor types but exhibits varying ΔPSI magnitudes (Figure 7B). Only bladder carcinoma (BLCA), B-cell lymphoma (DLBC), LAML acute myeloid leukemia (LAML), lung adenocarcinoma (LUAD), and testicular tumors (TGCT) showed a decrease in inclusion greater than 10% ΔPSI, whereas the remaining tumor types showed a significant splicing change of less than 10% between MYC-active and -inactive tumors. Conversely, increased inclusion was observed in kidney chromophobe (KICH) and uveal melanoma (UVM) in MYC-active tumors (Figure 7B). These findings indicate that many AS events associated with MYC-activity in breast cancer are also found in other tumor types.

**Figure 7.**
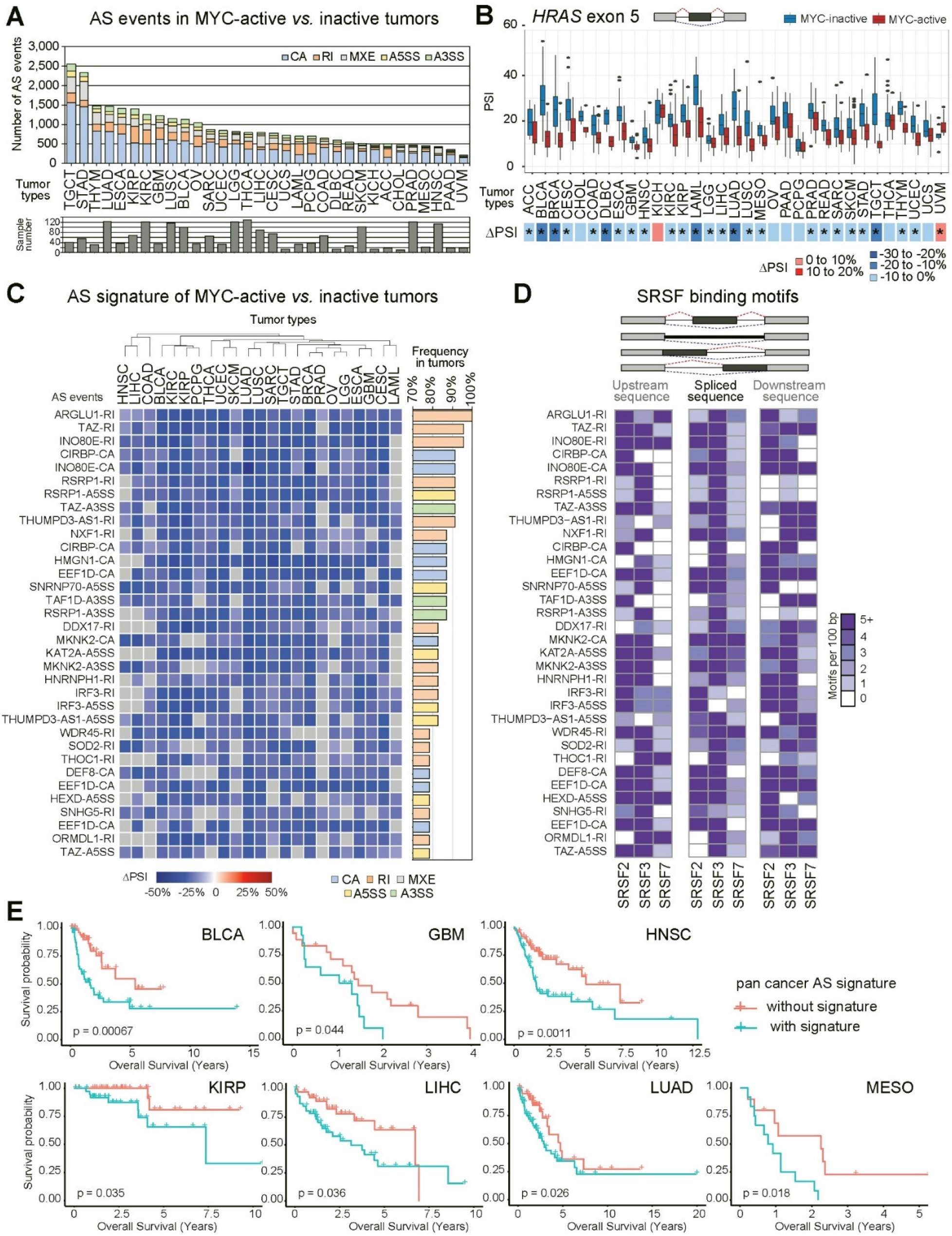
Pan-cancer splicing signature correlates with MYC activity and worse patient survival. **(A)** Significant AS events detected in MYC-active *vs.* -inactive TCGA tumors (ΔPSI>|10%|, FDR<0.05), shown across 33 TCGA tumor types (see tumor names in Table S2E), color-coded per AS event type. The number of samples in each tumor type is indicated. **(B)** Percent-spliced-in (PSI) values of an *HRAS* exon 5 splicing event detected in MYC-active (red) *vs.* - inactive (blue) tumors, shown per tumor type (median±interquartile range). ΔPSI (MYC-active *vs.* -inactive tumors) are shown as a heatmap below per tumor type. Significant ΔPSI changes (FDR<0.05) are indicated by an asterisk. **(C)** Differential splicing of 34 pan-cancer events in MYC-active *vs.* -inactive tumors across 75% of 23 TCGA tumor types (n>15 per group). Each column represents one tumor type and each row represents the differential percent-spliced-in (ΔPSI) value of a specific AS event in MYC-active *vs.* -inactive tumors, shown per tumor type. AS event type and gene name are shown. The frequency of each event across all 23 tumor types is plotted on the right, color-coded per AS event type. **(D)** Frequency per 100bp of predicted binding motifs for SRSF2, SRSF3, or SRSF7 in pan-cancer MYC-regulated AS events, in the upstream sequences (left panel), spliced sequence (middle panel), and downstream sequence (right panel). Individual motifs were scored as in Fig. 4A and then summed for each SR protein. **(E)** Kaplan-Meier plots for overall survival of cancer patients with or without the pan-cancer MYC AS events signature, shown per TCGA tumor type (log-rank test *P*-values). See also Figure S8 and Table S5.

We systematically filtered for shared AS events detected in ≥25% of all MYC-active tumors at |ΔPSI|≥10% and FDR<0.05 cutoffs, and identified 695 AS events across multiple TCGA tumor types (Figure S8A). These spliced genes were most enriched in pathways involved in RNA splicing and processing (Figure S8B). To further identify AS changes shared by the majority of cancer patients, we focused on 23 tumor types that had RNA-seq data for ≥15 samples in both the MYC-active and -inactive groups. We identified 34 pan-cancer AS events differentially spliced in ≥75% of these 23 tumor types, which were all associated with a decrease in PSI value in MYC-active tumors, and enriched for intron retention events (Figure 7C). Spliced genes included cancer-associated kinases (*MKNK2*), genes implicated in mRNA processing (*ARGLU1*, *CIRBP*, *SNRNP70*, *NXF1*, *DDX17*, *RSRP1*, *HNRNPH1, THOC1*), transcriptional regulation and/or chromatin remodeling (*ARGLU1, TAZ*, *INO80E*, *HMGN1*, *KAT2A*, *TAF1D, ZGPAT*), translational regulation (*EEF1D*), autophagy (*WDR45*), or metabolism (*HEXD*, *SOD2*, *ORMDL1*) (Figure 7C). Intron retention events often introduce a premature termination codon; decreased intron retention as observed in MYC-active tumors would therefore be predicted to lead to increased protein expression. For example, decreased *ARGLU1* intron retention, which is detected in all 23 MYC-active tumors, is predicted to increase ARGLU1 expression. *ARGLU1* is a transcriptional coactivator and splicing modulator that has been previously shown to be required for estrogen receptor-mediated gene transcription and breast cancer cell growth (Zhang et al., 2011). Decreased *ARGLU1* intron retention could therefore lead to increased cell growth, as well as anchorage-dependent and -independent colony formation of breast cancer cells as described (Zhang et al., 2011). SRSF3 and SRSF7 have been shown in mouse to directly bind to the *ARGLU1* retained intron (Brugiolo et al., 2017). On the other hand, intron retention in *NXF1*, encoding the nuclear export factor also known as ALY/REF, produces a truncated protein that lacks LRR, TAPC and NTF2 domains important for nuclear export, but is an important co-factor for the full-length isoform (Li et al., 2016). Interestingly, both SRSF3 and SRSF7 have been shown to promote NXF1 recruitment to mRNA, thus coupling AS and polyadenylation to NXF1-mediated mRNA export to control the cytoplasmic abundance of transcripts with alternative 3′ ends (Huang et al., 2003; Muller-McNicoll et al., 2016). Similarly, the use of the alternative splice site in *KAT2A* is predicted to produce a truncated protein that lacks the acetyltransferase and bromodomains. Four of the pan-cancer splicing events (*ARGLU1*, *NXF1, KAT2A,* and *WDR45*) were validated by RT-PCR in MCF-10A MYC-ER cells with MYC activation (Figure 3I, S3L), suggesting they are indeed MYC-regulated.

To predict which pan-cancer AS events are regulated by pan-cancer SF-module hub genes, SRSF2, SRSF3 or SRSF7, we evaluated whether the alternative sequences and flanking exons contain binding motifs for these three SFs. All 34 pan-cancer AS events contained motifs for *SRSF2*, *SRSF3*, and/or *SRSF7* (Figure 7D), with the majority containing motifs for all three SFs, suggesting these SR proteins may be responsible for regulating their splicing.

Finally, we scored the pan-cancer splicing signature across 23 TCGA tumor types, ranking tumors based on the inclusion or skipping patterns of each of the 34 pan-cancer AS events from Figure 7C. Patients with the pan-cancer AS signature had worse overall survival (Liu et al., 2018) compared to patients without the signature across multiple tumors types, *i.e.*, bladder carcinoma (BLCA), glioblastoma (GBM), head-neck squamous cell carcinoma (HNSC), kidney renal papillary cell carcinoma (KIRP), liver hepatocellular carcinoma (LIHC), lung adenocarcinoma (LUAD), and mesothelioma (MESO) (Figure 7E). These 34 AS events therefore represent a pan-cancer MYC-active signature of potential clinical utility, and indicate shared splicing regulatory networks across multiple tumor types.

## DISCUSSION

Previous studies have shown that MYC can regulate individual SFs that cooperate with MYC to promote tumorigenesis, such as *SRSF1* and *PRMT5* (Anczukow et al., 2012; Das et al., 2012; Koh et al., 2015). In addition, MYC has been associated with the expression of certain cassette exon or intron retention events in prostate, lymphoma, and breast cancers (Hsu et al., 2015; Koh et al., 2015; Phillips et al., 2020). However, a global understanding of how, and to what extent, MYC regulates splicing mechanisms has been missing. This study provides a comprehensive view of MYC-regulated splicing across 33 tumor types.

### Co-expression of MYC-regulated splicing factors in human tumors

Our study identified 18 SF co-expression modules in two independent breast cancer cohorts, six of which had a high correlation with MYC activity and were preserved to varying degrees in other tumor types. We validated the MYC-induced co-expression of SFs in cell models. We demonstrated that SR protein family members *SRSF2*, *SRSF3*, and *SRSF7* are co-expressed hub genes of a pan-cancer module in MYC-active human cancers. This pan-cancer SF-module controls cell invasion and induces the expression of a subset of MYC-regulated alternatively spliced isoforms, some of which have been previously associated with cell invasion, including *ENAH* and *PUM2* (Balsamo et al., 2016; Wang et al., 2019b; Zhang et al., 2019). Our findings suggest that coordinated expression of *SRSF2*, *SRSF3* and *SRSF7* under MYC-active conditions plays a role in MYC-driven tumorigenesis. *SRSF2*, *SRSF3* and *SRSF7* have been previously implicated in cancer individually and are upregulated in several tumor types compared to normal tissue (Park et al., 2019; Urbanski et al., 2018). *SRSF2* is commonly mutated in hematopoietic malignancies including myelodysplastic syndrome (MDS), chronic myelomonocytic leukemia (CMML) and acute myeloid leukemia (AML), and upregulation of *SRSF2* has also been reported in solid tumors such as breast and hepatocellular carcinoma (Luo et al., 2017; Park et al., 2019). *SRSF3* is upregulated in many tumor types including breast, brain, ovarian, stomach, bladder, colon, and liver cancers, and is emerging as a potential oncogene (Jia et al., 2010; Song et al., 2019; Urbanski et al., 2018). Although less is known about the role of *SRSF7* in cancer, it has been implicated in lung and colon cancer (Fu and Wang, 2018; Saijo et al., 2016; Urbanski et al., 2018). Interestingly, overexpression of *SRSF2* or *SRSF3* individually in non-transformed mammary epithelial cells did not result in an oncogenic phenotype (Park et al., 2019), suggesting that co-expression of these SR proteins might be required to promote tumor formation and maintenance.

We found several other SFs previously associated with MYC clustering in SF-modules in breast tumors. For example, *BUD31* and *hnRNPA1*, two SFs important for MYC-driven tumors (David et al., 2010; Hsu et al., 2015), were found in SF-modules #4 and #5 respectively, suggesting these modules are also likely important for MYC oncogenic activity. These modules were preserved in 25 and 32 of the 33 tested TCGA tumor types, respectively. *PRMT5*, despite being a known MYC target, was not found in any of the breast SF modules, suggesting its role in MYC-driven tumors may be specific to other tumor types such as lymphoma (Koh et al., 2015). We also found several SF-modules to be highly preserved in certain tumor types but not in others, suggesting tumor-specific biological roles. For example, SF-module #4 is highly preserved in stomach (STAD) but not in ovarian cancer (OV). The top gene in SF-module #4, *SNRPA1*, is a component of the U1 snRNP and has previously been implicated in gastric tumor growth (Dou et al., 2018). However, the preservation of a module *per se* is not sufficient to define that the expression of that module is under MYC-regulated control, and thus it is necessary to examine whether MYC activity correlates with module expression in a given tumor type. While four SF-modules with the highest correlation with MYC activity in breast tumors maintained high correlation in most other tumor types, two SF-modules (#4 and #6) showed variable correlation in some tumors. For instance, module #4 had a weak correlation with MYC activity in pheochromocytoma (PCPG) and diffuse large B-cell lymphoma (DLBC) and even had a negative correlation in uveal melanoma (UVM). This suggests that, while some SF-modules may be important for MYC-driven oncogenesis in multiple tumor types, other modules may be under the control of MYC in a more tumor type-specific manner. Indeed, tissue-specific roles of MYC have been reported, and inactivation of MYC in different tumor types results in varying outcomes such as apoptosis or differentiation (Gabay et al., 2014; Kress et al., 2015)..

While most SF-modules positively correlated with MYC-activity in TCGA breast tumors, SF-module #18 had a strong negative correlation, and, concordantly, genes within this module were significantly downregulated upon MYC activation in MCF-10A cells. Although MYC is commonly thought of as a transcription activator, MYC can induce transcriptional repression of select genes (Kress et al., 2015). Two of the top genes in SF-module #18 are CLK1 and CLK4, part of the CDC2-like family of kinases that are responsible for phosphorylation of SR-proteins (Bates et al., 2017). CLK inhibitors are currently under investigation as a potential therapeutic option for tumors, including MYC active tumors, and *MYC* amplification has been associated with increased sensitivity to the CLK inhibitor T-025 (Iwai et al., 2018). However, the negative correlation between MYC activity and module #18 may be cohort-specific, as it was not maintained in SCAN-B breast tumors. Therefore, while here we focused on the function of SF-module #3 which exhibits a striking pan-cancer correlation with MYC activity, further investigation into other MYC-induced SF-modules is likely to shed important light into pathogenesis and treatment of MYC-driven tumors.

### Cancer-associated role of the pan-cancer SF-module hub genes

Co-expression of pan-cancer SF-module #3 hub genes impacts patterns of a subset of AS events that are also detected in MYC-active tumors and cell models. Studies from our lab and others have previously shown *SRSF2*, *SRSF3*, and *SRSF7* have both distinct and overlapping AS targets, yet all prior studies examined the effects of an individual SF (Bradley et al., 2015; Long and Caceres, 2009; Park et al., 2019; Van Nostrand et al., 2020a). We demonstrate here that co-expression of these three SFs can impact MYC-driven splicing via a combination of mechanisms. First, each SF has distinct AS targets (*e.g.*, *NCOR1, PUM2*), resulting in expression of different alternative isoforms that together might promote tumor invasion. Second, SFs share targets to promote either skipping or inclusion of the same exon (*e.g.*, *HRAS*); this cooperation could lead to even higher levels of that oncogenic isoform. This could occur by multiple SFs binding the same transcript increasing the likelihood of splicing that exon, or increased SF expression could mean there is more SF present to bind transcripts and promote splicing. Third, SFs could have opposing effects on shared AS targets (*e.g.*, *BAG6*, *EHBP1, RAB25*). For instance, one SF could outcompete the other, via differences in binding affinity or number and/or location of binding sites. Our study showcases examples of MYC-regulated AS events that fall into these distinct categories, suggesting complex regulatory consequences of *SRSF2*, *SRSF3*, and *SRSF7* co-expression. However, the precise AS events that are required to drive tumorigenesis remain to be determined. Based on previous work from our lab and others, we anticipate that a combination of AS events is required to drive cancer-associated phenotypes, and modulation of a single AS event is unlikely to impact all the phenotypes associated with MYC-activation.

SR proteins are known to exhibit splicing-independent functions (Caceres et al., 1998; Huang and Steitz, 2001; Maslon et al., 2014; Muller-McNicoll et al., 2016; Sanford et al., 2004; Zhang and Krainer, 2004). We do not rule out that the roles of *SRSF2*, *SRSF3*, and *SRSF7* in other RNA processing steps, such as mRNA export, stability, and translation, may influence the phenotype of MYC-active tumors. Additionally, SFs have been shown to form extensive cross-regulatory networks via coupling of AS and nonsense-mediated mRNA decay (NMD) to regulate their expression via splicing of a non-coding poison exon in the SF gene (Leclair et al., 2020). In fact, *SRSF3* and *SRSF7* induce inclusion of each other’s poison exon, leading to decreased protein levels, whereas they induce skipping of the poison exon in *SRSF2* and lead to increased SRSF2 levels (Leclair et al., 2020). We posit that MYC’s regulatory effect overcomes SR protein negative cross-regulation, and that MYC’s control of this poison exon-based regulatory network is a critical step in MYC-driven transformation and tumor maintenance.

### MYC-active splicing signature in human tumors

Genes alternatively spliced in MYC-active breast cancer tumors were enriched in pathways involved in RNA splicing and processing, similar to findings in prostate cancer where MYC-correlated cassette exons were enriched in RNA binding protein genes (Phillips et al., 2020). The significance of this observation remains to be determined, but Phillips *et al*. speculate that MYC-correlated alternative exons in RNA binding proteins contain premature termination codons and could therefore be involved in AS-NMD regulation of SFs. Our lab has previously shown that inclusion of non-coding poison exons in SFs is significantly lower in tumors compared to normal samples (Leclair et al., 2020). Additionally, splicing of RNA binding proteins could lead to expression of isoforms that indirectly promote expression of oncogenic isoforms via splicing-independent mechanisms.

Our analysis revealed that all MYC-active tumors display differentially spliced events compared to MYC-inactive tumors. Given that MYC can have tissue-specific effects, investigation into the spliced isoforms within each tumor type is required to elucidate their role in MYC-driven tumor maintenance. However, we also identify a MYC-active splicing signature shared across >75% of 23 different tumor types, which provides a pan-cancer tumor classifier of MYC status, and can be used to predict prognosis and to identify patients most likely to benefit from splicing-modulating therapies. The use of splicing signatures to predict prognosis has already been proposed for a variety of tumor types individually, including breast, hepatocellular, and renal (Duan and Zhang, 2020; Wang et al., 2020b; Zhang et al., 2009). Our signature is remarkable in that it correlates with lower survival in several tumor types. MYC status in tumors is typically classified based on *MYC* copy number and/or expression, but these measures do not give a complete picture of MYC activity due to their lack of consideration of important regulators of MYC such as binding partners and phosphorylation status (Conacci-Sorrell et al., 2014; Hann, 2006). The use of alternative measures of MYC activity, such as a splicing signature, may prove to be a superior method of classifying tumors leading to better prediction of prognosis and/or treatment responses.

### Targeting splicing in MYC-driven cancer

The importance of co-expression of several SFs for invasion of MYC-driven tumors means that methods targeting these SFs or their downstream splicing targets may be a potential avenue for treatment. It may not be enough to target individual genes or SFs, but instead necessary to consider how genes cooperate. In particular, the degree to which the genes in pan-cancer SF-module #3 must be modulated to provide therapeutic benefit remains to be determined.

MYC-active tumors have been shown to be especially susceptible to splicing inhibition or disruption of the splicing machinery, for example, using spliceosomal inhibitors or SF-targeting shRNAs (Bowling et al., 2021; Hsu et al., 2015; Hubert et al., 2013; Iwai et al., 2018; Koh et al., 2015), suggesting MYC confers an increased dependence on the splicing machinery (Hsu et al., 2015). Alternatively, splicing inhibition in MYC-active breast cancer can result in the accumulation of double-stranded RNA, which triggers an antiviral response and apoptosis (Bowling et al., 2021). Splicing disruption with splicing-targeted therapeutics, including small molecule inhibitors or RNA-based splice-switching approaches, can lead to alterations in SF activity and/or alternative isoform expression (Corrionero et al., 2011; Teng et al., 2017; Vigevani et al., 2017). Whether these alterations occur in MYC-regulated SFs or alternative isoforms remains to be determined.

An example that demonstrates how MYC-regulated splicing may be of particular relevance to splicing-targeted therapy is with inhibitors of the SF PRMT5, such as EPZ015666 (also known as GSK3235025). PRMT5 modifies snRNP-associated Sm proteins and is important for MYC-driven tumorigenesis in lymphoma (Koh et al., 2015). Are PRMT5 inhibitors most efficacious in tumors with high MYC activity? Do these inhibitors have effects on MYC-regulated splicing and are such effects either a cause for, or a correlate of, selective sensitivity? If so, should clinical trials seek to characterize MYC activity and specifically MYC-regulated splicing in tumors prior to treatment? These questions will also be important to address for other emerging splicing-modulating therapies in MYC-driven tumors for several reasons. First, if MYC-regulated spliced isoforms regulate or associate with tumor sensitivity to therapies, it will be critical to consider the splicing profile of tumors in patients enrolled in clinical trials. If a tumor does not express a specific isoform, treatment with a therapy that targets that isoform may not be effective. Second, uncovering which aspects of MYC-regulated splicing are crucial could lead to new strategies of targeting these regulators or isoforms. For example, the development of splice-switching antisense oligonucleotides (ASOs) which can target individual MYC-regulated splicing events might have fewer side effects than broader splicing inhibition approaches.

Identification of cancer-specific and pan-cancer mutational patterns have been critical in improving our understanding of what drives tumor growth and progression, and has also led to the development of cancer therapies and biomarkers for prognosis (Martinez-Jimenez et al., 2020). We expect that a fuller characterization of pan-cancer and tumor-specific molecular alterations, which, as we show here, include co-expressed SFs and AS events regulated by MYC, will provide additional benefits for both furthering our understanding of tumor biology and the development of therapies.

## LIMITATIONS

Although we detected a large number of MYC-regulated AS events in more than one dataset, *e.g.*, TCGA, HCC1806, or MCF-10A, some events were either changing in opposing directions or were not detected as being MYC-activated in one of the cell lines. These differences are likely explained at least in part by technical and biological differences. First, differences between the quality of the RNA-seq datasets used in this study, and in particular the smaller library sizes, shorter read length, and lower overall quality of the TCGA project (∼30-40 million 50bp reads) compared to our data from cell models (>100 million 150 bp reads), likely limits our ability to detect TCGA AS events in genes expressed at lower levels. MCF-10A cells are non-transformed mammary epithelial cells and therefore some non-overlapping effects of MYC between this system and established tumors are to be expected. Additionally, MYC in MCF-10A was induced over a short period, whereas TCGA MYC-active breast tumors have chronic MYC activation. It is possible that events induced initially by MYC differ from those expressed and regulated by MYC after sustained MYC activation. Moreover, although HCC1806 was classified as a MYC-inactive breast cell line according to our analysis, these cells contain other mutations and alterations that may influence MYC activity upon overexpression. Additionally, the baseline expression of certain isoforms in these cells is elevated and therefore MYC activation did not induce significant further changes in their splicing. Finally, although the overexpression levels in cell lines are comparable to those in MYC-active breast tumors, where SRSF2, SRSF3, and SRSF7 are upregulated by 1.5-2 fold, SF-module #3 overexpression in HCC1806 cells is milder than in MCF-10A cells, likely due to the fact that higher expression of multiple SR proteins simultaneously can be toxic to cancer cells.

## ACKNOWLEDGMENTS

We thank Jeffrey Chuang, Brenton Graveley, Stefan Pinter, Gordon Carmichael, and Ching Lau for helpful discussions, Taneli Helenius for extensive comments and edits on the manuscript, Senthil Muthuswamy for the gift of MCF-10A and MCF10A MYC.ER cells, and Edison Liu for HCC1806 cells. We thank Christina Chatzipantsiou and Anne Deslattes Mays for assistance with implementing our splicing analysis pipeline in the cloud. We acknowledge assistance from Microscopy, Single Cell Biology Laboratory and Genome Technologies Sequencing Services at The Jackson Laboratory (JAX). This work was supported by NIH grants R00CA178206, R01CA248317 and R01GM138541 to OA, T32AG062409A to BLA, and NCI P30CA034196 to JAX. We acknowledge the use of data generated by TCGA, managed by NCI and NHGRI, downloaded using ISB Cancer Genome Cloud and processed using Google Cloud Platform Research Credit award. The content is solely the responsibility of the authors and does not necessarily represent NIH official views. The results published here are in part based upon data generated by the TCGA Research Network: https://www.cancer.gov/tcga and CCLE: https://portals.broadinstitute.org/ccle.

## AUTHOR CONTRIBUTIONS

LU and OA designed the study, analyzed data, and wrote the paper. LU, MB, SHP, NL, and BLA conducted experiments. LU, BLA, PP and SKS implemented the cloud-based analysis pipeline. LU and BLA performed bioinformatics analyses. OA supervised the study. All authors discussed the results and manuscript.

## DECLARATION OF INTERESTS

None.

## KEY RESOURCES TABLE

| Reagent or resource | Source | Identifier |
| --- | --- | --- |
| **Antibodies** |  |  |
| Anti-SRSF1 mouse | CSHL antibody facility | Ak96322 |
| Anti-TRA2B rabbit | Abcam | #Ab31353 |
| Anti-Beta-Catenin | ThermoFishe | #71-2700 |
| Phalloidin-A647 | ThermoFisher | #A22287 |
| Alexa Fluor 568 anti-mouse | Invitrogen | #A-11031; RRID: AB_144696 |
| Alexa Fluor 488 anti-rabbit | Invitrogen | #A-11034; RRID: AB_2576217 |
| IRDye 800CW Goat anti-Rabbit IgG (H + L) | Li-Cor | #926-32211; RRID:AB_621843 |
| IRDye 680 Goat anti-Mouse IgG (H + L) | Li-Cor | #926-68070; RRID:AB_10956588 |
| **Chemicals, peptides, and recombinant proteins** |  |  |
| Hoechst | Invitrogen | #62249 |
| Calcein, AM | Invitrogen | #C3099 |
| Matrigel Growth Factor Reduced Phenol-free | BD/Corning | #356238 |
| SYBRSafe | Invitrogen | #S33102 |
| Superscript III reverse transcriptase | Invitrogen | #18080044 |
| Trypsin-EDTA 0.25% | Gibco | #25200056 |
| **Critical Commercial Assays** |  |  |
| RNAeasy kit | Qiagen | #74106 |
| Lipofectamine 2000 | Invitrogen | #11668019 |
| Phusion flash high fidelity master mix | Thermo Fisher | #F548L |
| Phusion hot start II DNA polymerase | Thermo Fisher | #F549L |
| iTaq Universal SYBR green Supermix | BioRad | #1725122 |
| TrueSeq stranded mRNA kit with polyA selection | Illumina | #20020594 |
| Cell Recovery Solution | BD/Corning | #354253 |
| **Deposited Data** |  |  |
| RNA-seq MCF-10A MYC-ER | This paper | pending |
| RNA-seq HCC1806 | This paper | pending |
| **Experimental Models: Cell lines** |  |  |
| MCF-10A | Muthuswamy lab (BIDMC) | N/A |
| MCF-10A MYC-ER | Muthuswamy lab (BIDMC) | N/A |
| HEK293T | ATCC | CRL-3216 |
| 293GPG | Muthuswamy lab (BIDMC) | N/A |
| HCC1806 | Liu Lab (JAX) | N/A |
| MDA-MB231 GFP-luciferase<br>rTTA3-Puro | (Park et al., 2019) | N/A |
| **Recombinant DNA** |  |  |
| pBABE-Puromycin | AdGene | #1764 |
| pWZL-Hygromycin | Lowe Lab (MSKCC) | N/A |
| pWZL-MYC-Hygro | Lowe Lab (MSKCC) | N/A |
| PWZL-T7-SRSF2-Hygro | (Park et al., 2019) |  |
| pBABE-T7-SRSF3-Puro | This paper | N/A |
| PWZL-HA-SRSF7-PuroG418 | This paper | N/A |
| TRMPV-Neo | Zuber et al. 2011 |  |
| TRMPV-Neo shRNA control | (Park et al., 2019) |  |
| TRMPV-Neo shRNA SRSF2 | This paper | N/A |
| TRMPV-Neo shRNA SRSF3 | This paper | N/A |
| TRMPV-Neo shRNA SRSF7 | This paper | N/A |
| pCI-neo | Krainer Lab (CSHL) | (Hua et al., 2007) |
| pCI-neo-HA-empty vector | (Leclair et al., 2020) | N/A |
| pCI-neo-HA-SRSF2 | (Leclair et al., 2020) | N/A |
| pCI-neo-HA-SRSF3 | (Leclair et al., 2020) | N/A |
| pCI-neo-HA-SRSF7 | (Leclair et al., 2020) | N/A |
| **Sequence-based reagents** |  |  |
| RT-PCR primer sequences | IDT | Table S6 |
| SLIC cloning primers? | IDT | Table S6 |
| **Software and Algorithms** |  |  |
| STAR (v.2.7.3a) | (Dobin et al., 2013) | https://github.com/alexdobin/STAR |
| Stringtie (v.2.0.6) | (Pertea et al., 2015) | https://ccb.jhu.edu/software/stringtie/ |
| Trimmomatic (v.0.39) | (Bolger et al., 2014) | http://www.usadellab.org/cms/?page=trimmomatic |
| DESeq2 | (Love et al., 2014) | N/A |
| rMATS (v.4.0.2) | (Phillips et al., 2020; Shen et al., 2014) | https://github.com/Xinglab/rmats-turbo |
| splicing-pipelines-nf v1.0 | This paper | https://github.com/TheJacksonLaboratory/splicing-pipelines-nf |
| WGCNA | (Langfelder and Horvath, 2008) | N/A |
| NetRep | (Ritchie et al., 2016) | https://github.com/sritchie73/NetRep |
| RStudio | N/A | N/A |
| GraphPad Prism | GraphPad | N/A |
| Harmony High-Content Imaging and Analysis Software | Perkin Elmer | N/A |
| ImageJ Digital Image Processing Software | (Schneider et al., 2012) | https://imagej.nih.gov/ij/ |
| RBPmap | (Paz et al., 2014) | http://rbpmap.technion.ac.il/ |
| Photoshop and Illustrator CC2019 | Adobe | N/A |
| TCGAbiolinks | (Colaprico et al., 2016; Mounir et al., 2019; Silva et al., 2016) | N/A |
| ChemiDoc MP Imaging System | BioRad | N/A |
| QuantStudio Real-Time software | ThermoFisher | N/A |

## EXPERIMENTAL MODEL AND SUBJECT DETAILS

### Human cell lines

MCF-10A and MCF-10A MYC-ER cells were a gift from Senthil K. Muthuswamy (Beth Israel Deaconess Medical Center) and were maintained in DMEM/F12 (Gibco) supplemented with 5% horse serum (GIBCO), 1% penicillin streptomycin (Sigma), 20 ng/mL EGF (Peprotech), 2 ug/mL hydrocortisone 0.5 ug/mL (Sigma), 100 ng/mL cholera toxin (Sigma), and 10 ug/mL insulin (Sigma) (Debnath et al., 2003).

HCC1806 cells, a gift from Edison Liu (Jackson Laboratory), were maintained in DMEM (Gibco) supplemented with 15% FBS and 1% penicillin streptomycin (Sigma).

HEK293T cells (ATCC), were maintained in DMEM (Gibco) supplemented with 10% FBS, 1% penicillin streptomycin (Sigma).

MDA-MB231 GFP-luciferase rTTA3-puro (Park et al., 2019) were maintained in DMEM (Gibco) supplemented with 20% FBS, 1% penicillin streptomycin (Sigma).

All cell lines were grown at 37°C under a humidified atmosphere with 5% CO_2_. Cells routinely tested negative for mycoplasma using the MycoAlert™ Mycoplasma Detection Kit (Lonza). Cell aliquots from early passages were used.

## METHOD DETAILS

### Plasmids

The pBABE-T7-SRSF3-Puro plasmid was previously described (Park et al., 2019). The pWZL-T7-SRSF2-Puro plasmid was created by subcloning T7-SRSF2 cDNA from a pWZL-T7-SRSF2-Hygro (Park et al., 2019) into a pBABE-Puro plasmid (a gift from S. Lowe, Memorial Sloan Kettering Cancer Center) using sequence and ligation independent cloning (SLIC). The pWZL-G418 plasmid was created by replacing the hygromycin resistance DNA sequence in pWZL-Hygro (a gift from S. Lowe, Memorial Sloan Kettering Cancer Center) by G418 resistance DNA sequence from TRMP-Neo (Zuber et al., 2011). The PWZL-HA-SRSF7-G418 plasmid was created by subcloning HA-SRSF7 cDNA from the pCI-neo-HA-SRSF7 plasmid (Leclair et al., 2020) into the pWZL-G418 plasmid using SLIC. The pBABE-HA-MYC-Hygro used here was a gift from S. Lowe (Memorial Sloan Kettering Cancer Center). Corresponding empty vector plasmids, pBABE-Puro, pWZL-Hygro, and PWZL-G418 were used as control.

The pCI-neo-HA-SR-CDS plasmids were previously described (Leclair et al., 2020), and contain the coding sequence (CDS) of each human SR protein along with an sequence encoding the HA-tag cloned into a pCI-neo mammalian expression plasmid (Hua et al., 2007).

Short hairpins targeting SR proteins were designed using SplashRNA (Pelossof et al., 2017) and subcloned into a TRMPV-Neo plasmids (Zuber et al. 2011) as previously described (Park et al., 2019).

SLIC cloning primers are shown in Table S6B.

### Generating stable cell lines

HCC1806 expressing T7-tagged SRSF2, T7-tagged SRSF3, and HA-tagged SRSF7 cDNA, alone or in combination, as well as corresponding empty vector controls and HA-tagged MYC, were generated by retroviral transduction as described (Anczukow et al., 2012; Park et al., 2019) via successive rounds of infection and selection. Virus was produced by transfection 15 ug of plasmid in 293GPG cells (a gift from S. Muthuswamy, BIDMC) using Lipofectamine 3000 (Invitrogen) per manufacturer instructions, along with helper packaging plasmids. Virus collected after transfection was concentrated by adding 0.09x PBS, 0.3M NaCl and 8.5% PEG6000, the solution was gently mixed at 4°C overnight, and the viral concentrate was spun for 15 min at 7000g and the supernatant gently removed. Viral particles were resuspended with appropriate volume of OptiMem (Invitrogen) and either immediately used or frozen at –80C. Stable cells were selected using 1-2 µg/ml puromycin (Gibco), 30-50 µg/ml hygromycin (Invitrogen), or 500 µg/ml Geneticin (G418) (GoldBio) until non-infected control cells were all dead. Cell lines were then maintained in media supplemented with 0.5 µg/ml puromycin, 15 µg/ml hygromycin and 500 µg/ml Geneticin (G418) to reduce excessive antibiotic stress on the cell lines.

MDA-MB231-luciferase-GFP-rTTA3 and HCC1806-MYC -rTTA3 lines expressing SF and control-shRNA-TRMPV-Neo were generated as retroviral transduction and selection as described (Park et al., 2019). Multiple shRNAs were tested for each target, and the most efficient RNA was selected. shRNA sequences are shown in Table S6E.

### SR protein co-expression in HEK293T cells

HEK293T cells were reverse transfected in 24 well plates at a seeding density of 600,000 cells/mL using Lipofectamine 3000 (Invitrogen) according to manufacturer’s instructions. At the time of transfection 750ng of total plasmid was diluted into 100uL OptiMem (Invitrogen). For individual SR protein transfections this included 250 ng of pCI-neo-HA-SR with 500ng of pCI-neo-HA-empty vector, double SR protein transfections included 250ng of each SR encoding plasmid with 250ng of the control plasmid, triple SR protein transfections included 250ng of each SR encoding plasmid, and the control well was 750ng of HA-empty vector. 48h after transfection the cells were collected by lifting with 2mM EDTA.

### 2D Transwell migration assays

HCC1806 cells were starved in serum-free media for 4h before seeding 200,000 cells in serum free media on top of an 8-μm PET membrane transwell (BD-Biosciences) in a 24-well format and allowed to migtrate into the lower compartment containing media supplemented with 15% of FBS for 24 hours. After 24h the cells on the top of the filter were removed by scraping with a que-tip and remaining cells under the filter were fixed using 5% Formalin (Sigma), then permeabilized with 0.5% Triton X-100 (Sigma) and stained with DAPI (Invitrogen). DAPI-positive cells were imaged using Zoe Fluorescent Cell Imager (Bio-Rad).

### 2D Cell proliferation assays

HCC1806 or MDA-MB231cells were plated in 96-well plate at 5,000 cells per well. For MDA-MB231 cells media was supplemented with 2 μg/ml of doxycycline (Sigma) or mock. Cell number was inferred via luminescence measurement using the Cell Titer Glo (Promega) assay per manufacturer instructions at day 1, 3, 5 and 7 for HCC1806, or day 1, 2, 3, 4 for MDA-MB231, using a Synergy H1 microplate reader and imager. For each sample, relative luminescence was normalized to luminescence on day 1, for 3-4 biological replicates at each timepoint.

### 3D cell culture assays and imaging

For 3D culture assays, MCF-10A and MCF-10A MYC-ER cells were seeded at a density of 10,000 cells per well in triplicate on a 4-well glass chamber slide coated with Matrigel Growth Factor Reduced (BD Biosciences) as described (Debnath et al., 2003; Liberzon et al., 2015). Media was replaced at 72h. Starting on day 3, cells were treated with 1μM 4-hydroxy tamoxifen (4-OHT) (Sigma) for 48h, 24h, 16h, 8h, 4h, or 0h, prior to collection. All samples were collected at the same time on day 5.

For 3D-culture assays, HCC1806 cells were seeded at a density of 15,000 cells per well in a 48-well tissue culture plate coated with 125μl of Matrigel Growth Factor Reduced (BD Biosciences). Media was replaced every 72h and growth was monitored for 9 days. For HCC1806-MYC-OE-rTTA3-shRNA cells, media was supplemented with 2 μg/ml of doxycycline (Sigma) or mock. At days 5 or 9, organoids were treated for 15 minutes with Calcein AM (1μM final concentration, Invitrogen) and Hoechst (1x final concentration, Invitrogen) diluted in 1x PBS, and multiple fields and z-stacks were imaged for each well using the Opera Phenix High-Content Screening System (PerkinElmer). Maximal projection of all imaged fields and z-stacks were reconstructed using the Harmony High-Content Imaging and Analysis Software (PerkinElmer) and analyzed using ImageJ digital processing software (https://imagej.nih.gov/ij/), only structures with a total area bigger than 700 μm^2^ were considered in the analysis.

For 3D invasion assays, 15,000 HCC1806 cells were seeded on a 1:1 mix of collagen:matrigel (to reach a final concentration of Collagen of 1.6 mg/ml and pH was adjusted to be at ∼7.6) in a 48-well tissue culture plate. Media was replaced every 3-4 days. At day 9 organoids were treated for 15 minutes with Calcein AM (1μM final concentration, Invitrogen) in DMEM and multiple fields and z-stacks were imaged on the Opera-Phenix High-Content Screening System (PerkinElmer).

For 3D culture assays, MDA-MB231-rTTA3-shRNA cells were seeded at a density of 7,000 cells per well in triplicate for control and DOX-treated lines, on a 48-well plate coated with Matrigel Growth Factor Reduced (BD Biosciences). Media was replaced every 72h and supplemented with 2 μg/ml of doxycycline (ADD) or mock, and growth was monitored for 9 days. On day 9, organoids were treated for 15 minutes with 1μM final Calcein AM (Invitrogen) diluted in growth media. Multiple fields and z-stacks were imaged for each well using the Opera Phenix High-Content Screening System (PerkinElmer). Maximal projection was used to reconstruct representative Z-stack fluorescent confocal images of 25 fields with 30-35 Z-stack images spaced every 55 μm using the Harmony High-Content Imaging and Analysis Software (PerkinElmer). Maximal projection of all imaged fields and z-stacks was analyzed using ImageJ digital processing software (https://imagej.nih.gov/ij/) to calculate organoid area.

### 2D confocal imaging

HCC-1806 control, 3xSR, and MYC-OE cell lines were plated onto coverslips at low density. 24h later cells were fixed using 4% paraformaldehyde (Sigma), permeabilized, and stained with 5ug/ml of anti-β-catenin antibody (ThermoFisher) overnight at 4°C. Samples were then counterstained with 4 ug/ml Alexa-488 secondary antibody (Invitrogen), as well as 0.005 U/ul Alexa647-conjugated phalloidin (ThermoFisher), and 1 ug/ml DAPI (Invitrogen), and mounted onto slides using Prolong Gold Antifade reagent (Invitrogen). High resolution images were acquired using the 60x objective of an Dragonfly confocal microscope (Andor). Images represent the maximum intensity projection of an approximately 10 µm Z-stack encompassing the entirety of the cell. All post-acquisition image adjustments were made using ImageJ (https://imagej.nih.gov/ij/).

### RNA extraction

3D-grown MCF-10A cells were washed with PBS (1X) and the Matrigel was dissolved by incubating slides at 4°C in Cell Recovery Solution (BD Biosciences). 2D-grown HEK293T, HCC1806, or MDA-MB231 cells were harvested by scraping adherent cells in PBS once ∼90% confluence was reached. Total RNA was extracted using the RNeasy kit (Qiagen) including DNase I treatment per manufacturer instructions.

### RNA-sequencing

Barcoded RNA libraries were prepared starting with 1ug for MCF-10A and 500ng for HCC1806 cell lines of total RNA using the TrueSeq stranded mRNA kit with polyA selection (Illumina), and quantified using a Bioanalyzer DNA 1000 chip (Agilent). Libraries were sequenced as 150bp paired-end reads at 100-200 million reads per library on an Illumina HiSeq (MCF-10A) or NextSeq (HCC1806) instrument. For MCF-10A, equal amounts of 3 libraries were pooled per lane. For HCC1806, equal amounts of 18 libraries were pooled per lane. At least 3 independent biological samples were sequenced for each experimental condition, and run on separate lanes whenever feasible.

### Quantitative RT-PCR analysis

Total RNA from 3D-grown MCF-10A cells or 2D-grown MDA-MB231 or HCC1806 cells was extracted as described above. 1ug of total RNA was reverse-transcribed using Superscript III reverse transcriptase (Invitrogen). qPCR was used to amplify endogenous transcripts with SF specific primers (Table S6C) using cDNA corresponding to 5-20ng of total RNA. qPCR was performed with iTaq Universal SYBR green Supermix (Bio-Rad) in 384-well plates (Life Technologies) using a ViiA7 Real-Time PCR system (Life Technologies) per manufacturer instructions and analyzed with QuantStudio Real-Time software. SF-expression was normalized to housekeeping gene *GAPDH*.

### Semi-Quantitative RT-PCR analysis

Total RNA from 3D-grown MCF-10A cells was extracted as described above. 1ug of total RNA was reverse-transcribed using Superscript III reverse transcriptase (Invitrogen). Semi-quantitative PCR was used to amplify endogenous transcripts with with SF specific primers (Table S6C) using cDNA corresponding to 5-20ng of total RNA. Optimal PCR conditions were defined for each primer pair by testing amplification from 26-30 cycles to select semi-quantitative conditions. PCR products were separated by 2% agarose gel stained with SYBRSafe (Invitrogen), and bands were quantified with a ChemiDoc MP Imaging System (Bio-Rad). SF-expression was normalized to housekeeping gene *GAPDH*.

### RT-PCR splicing event validation

Total RNA from 3D-grown MCF-10A cells or 2D-grown HEK293T, MDA-MB231, or HCC1806 cells, was extracted as described above. 1ug of total RNA was reverse-transcribed using Superscript III reverse transcriptase (Invitrogen). Semi-quantitative PCR was used to amplify endogenous transcripts with primers that amplify both the included and skipped isoforms (Table S6D) using cDNA corresponding to 5-20ng of total RNA. Optimal PCR conditions were defined for each primer pair by testing amplification from 26-30 cycles to select semi-quantitative conditions. PCR products were separated by 2% agarose gel stained with SYBRSafe (Invitrogen), and bands were quantified with a ChemiDoc MP Imaging System (Bio-Rad). The ratio of each isoform was first normalized to the sum of the different isoforms, and changes were then expressed as the fold increase compared to the levels obtained for cells or organoids expressing the control vector.

### Western blot analysis

3D-grown MCF-10A and MCF-10A MYC-ER cells were washed with PBS (1X) and the Matrigel was dissolved by incubating slides at 4°C in Cell Recovery Solution (BD Biosciences). Cells were lysed in Laemmli buffer (50 mM Tris-HCl pH 6.2, 5% (v/v) β-mercaptoethanol, 10% (v/v) glycerol, 3% (w/v) SDS). Equal amounts of total protein were loaded on a stain-free 12% SDS-polyacrylamide gel (Biorad), transferred onto a nitrocellulose membrane (Millipore) and blocked in 5% (w/v) milk in Tween 20-TBST (50 mM Tris pH 7.5, 150 mM NaCl, 0.05% (v/v) Tween 20). Blots were incubated with TRA2β (Abcam), SRSF1 (CSHL), or β-catenin (ThermoFisher) primary antibodies. IR-Dye 680 anti-mouse or IR-Dye 800 anti-rabbit immunoglobulin G (IgG) secondary antibodies (LI-COR) were used for infrared detection and quantification with a ChemiDoc MP Imaging System (Bio-rad).

### Differential splicing analysis

Paired-end reads were preprocessed by trimming of low-quality regions by Trimmomatic (v. 0.39) (Bolger et al., 2014). Reads were then mapped to the human reference genome using STAR in 2-pass mode (v.2.7.3a) (Dobin et al., 2013) with the Gencode GRCh38 v.32 reference transcript annotation (Frankish et al., 2019). To include novel exons and introns in our analysis, we performed an annotation-guided transcriptome reconstruction and merged the resulting transcriptome (GTF) from each sample into one comprehensive transcript annotation using Stringtie (v.2.0.6) (Pertea et al., 2015). We utilized an in-house pipeline that implemented rMATS (v.4.0.2) (Shen et al., 2014) to detect splicing events using both splice junction read counts and alternatively spliced exon body counts (https://github.com/TheJacksonLaboratory/splicing-pipelines-nf v1.0). For each event, a percent spliced in (PSI) score was calculated. A ΔPSI is calculated for each event to compare the change in inclusion between MYC active and MYC inactive samples, such that a positive ΔPSI indicates increased inclusion in MYC active tumors whereas a negative ΔPSI indicates increased skipping. Differentially spliced events (DSEs) were filtered based on the following: i) ΔPSI=|mean PSI^case^ – mean PSI^control^ |≥0.1; and ii) FDR≤0.05; and iii) at least 5 reads (averaged across biological replicates) detected in both the control and case that support either exon skipping or exon inclusion, *i.e.*, (inclusion count ≥5 in either control OR case) AND (skipping count ≥5 in either control OR case).

To account for any 4-OHT-induced splicing in MCF-10A MYC-ER samples, we compared MCF-10A and MCF-10A MYC-ER differential splicing events at each time point, *i.e.,* MCF-10A 8h *vs.* MCF-10A MYC-ER 8h. 4-OHT induced differential splicing events were removed from downstream analysis if they met the following conditions: i) significant in both samples, and ii) had a ΔPSI with the same sign indicating a change in the same direction.

### Differential gene expression analysis

Preprocessing of reads and mapping steps were performed as described above using only the Gencode GRCh38 v.32 reference transcript annotation. A gene-level count matrix was generated using GTF files from Stringtie. Differential gene expression was performed using DESeq2 (Love et al., 2014). Genes with <10 total reads across samples were removed. A Wald test was used to calculate *P*-values, and Benjamini-Hochberg procedure was used to calculate corrected *P*-values. Differential genes were selected based on corrected *P*-value<0.05 and log_2_ fold change >0.5 or <-0.5.

To account for 4-OHT induced changes in expression in MCF-10A MYC-ER samples, genes that were significantly differentially expressed in the same direction in both MCF-10A control and MCF-10A MYC-ER sample were removed from further analysis.

### TCGA data gene expression and splicing analysis

Tumor and corresponding normal tissue samples were downloaded as bam files from the NCI Genomic Data Commons and processed on the used Lifebit’s Google Cloud Platform. Sample IDs are listed in Tables S1 and S5. Differential gene expression and splicing analysis were performed as described above.

### RBP protein motif analysis

SR protein RNA binding motifs compiled from literature (Table S6A) (Ajiro et al., 2016; Dominguez et al., 2018; Kim et al., 2015; Konigs et al., 2020; Paz et al., 2014; Ray et al., 2013) and from the RBPmap default list (Paz et al., 2014) were mapped to alternative exons and either surrounding 100 nt or upstream and downstream exons using the RBPmap webtool (http://rbpmap.technion.ac.il/) with default stringency and conservation filter cutoffs. The resulting motifs were visualized in the UCSC genome browser.

### Gene Ontology enrichment

Gene enrichment was performed using the *clusterProfiler* package (Yu et al., 2012). For AS enrichment analysis, we generated a gene list with all genes that contained a significant differential AS event detected using the reference GTF.

### Chromatin precipitation sequencing analysis

Peaks from ENCODE MYC ChIP-seq data from MCF-10A cells (ENCFF013XMV) (Consortium, 2012; Davis et al., 2018) were annotated using the *ChIPseeker* R package (Yu et al., 2015), with promoters defined as being within 1000 bp of the transcription start site.

### MYC-activity scoring in tumor samples

We implemented a MYC activity scoring system adapted from Jung et al. (Jung et al., 2017) using a list of known MYC target genes from MSigDB ‘Hallmark Gene Set MYC TARGETS V1’ (Liberzon et al., 2015). This scoring system was used to classify TCGA samples and for breast tumors from The Sweden Cancerome Analysis Network Breast Initiative (SCAN-B; GSE96058). Gene expression (normalized counts for TCGA samples and FPKM from SCAN-B tumors) were obtained, and all samples in each dataset were ranked based on their expression of MYC target genes in ascending order. The sum of all rank values (rank sum) was calculated for each sample and we then divided the rank sums by the average rank sum for the entire dataset. In order to compare TCGA normal breast with tumor samples, we classified all 1186 samples (113 normal adjacent, 1073 tumor) together. In order to classify MYC-active and -inactive TCGA tumors, normal adjacent samples were removed and MYC-activity scores were re-calculated. We then calculated MYC-activity z-scores for all tumors and MYC-active tumors were classified by having a z-score >1.5 whereas MYC-inactive tumors by z-score <-1.5. SCAN-B tumors were classified in the same manner. Since the number of tumors in other TCGA datasets is much less than that in breast, the threshold values for MYC-active z-scores were >1.2 and <-1.2 for all other TCGA tumor types, with the exception of CHOL which had a threshold of ±1.

### Weighted gene correlation network analysis (WGCNA)

SF co-expression analysis in TCGA breast tumors was performed with the *WGCNA* R package (Langfelder and Horvath, 2008, 2012) using log_2_ transformed gene expression normalized counts for 334 SFs. We used a signed network construction and a soft-thresholding power of 14, per WGCNA guidelines. To detect co-expression modules, we utilized the blockwiseModules function with a biweight mid-correlation, per WGCNA. Module eigengenes (1^st^ principal component) were calculated for each module and were used to calculate module membership (MM). MM is defined as the correlation of the module eigengene with gene expression profile and represents how close a gene is to that module. MM can be used to identify top genes, or hub genes, within a given module. For our analysis, we defined top genes for each module by having a MM>0.75.

### Co-expressed SF-module preservation analysis

Module preservation was determined for TCGA tumors and for breast tumors from the Sweden Cancerome Analysis Network Breast Initiative (SCAN-B, GSE96058). Gene expression data (FPKM) for 33 tumor types was obtained from TCGAbiolinks (Colaprico et al., 2016; Mounir et al., 2019; Silva et al., 2016). To evaluate preservation of the BRCA modules in other datasets, we utilized the R package NetRep (Ritchie et al., 2016). We first generated adjacency matrices as well as correlation matrices based on FPKM data for all 783 RPBs, and utilized the *modulePreservation* function. Since our modules were considered small, some of which had fewer than 10 nodes or genes, we only considered four parameters to determine if the module was significantly preserved as per NetRep guidelines: average edge weight (avg.weight), module coherence (coherence), average node contribution (avg.contrib), and average correlation coefficient (avg.cor.) Modules with a *P* <0.01 for at least three of the four parameters were considered significant.

### Survival analysis

Splicing inclusion values (PSI) for each of the 34 pan-cancer AS events were obtained as described above for each TCGA tumor type, and all samples in each dataset were ranked based on their PSI values. The sum of all rank values was calculated for each sample and we then divided the rank sums by the average rank sum for the entire dataset. We then calculated splicing z-scores for all tumors and tumors were classified based on their levels of inclusion of the signature using a z-score threshold >1.2. Survival data from TCGA was retrieved from (Liu et al., 2018) and Kaplan-Meier survival curves were plotted using R packages survival and survminer.

## QUANTIFICATION AND STATISTICAL ANALYSIS

Where appropriate, the data are presented as the mean±s.d., as indicated. Data points were compared using an unpaired two-tailed, Student t-test or two-tailed Mann-Whitney test, as indicated in the legends. For quantification of proliferation and apoptosis markers, a two-tailed Fisher test was used. *P*-values are indicated in the figure legends.

## DATA AND CODE AVAILABILITY

RNA-sequencing data has been deposited on GEO as GSE181968 for MCF-10A MYC-ER experiments, and GSE181956 for HCC1806 experiments.

RNA-sequencing data from TCGA tumors (The Cancer Genome Atlas Network, 2012) is available via ISB-CGC cloud. Sample IDs are listed in Tables S1 and S5.

ChIP-seq datasets (ENCFF013XMV) are available from https://www.encodeproject.org/.

Gene expression data for breast tumors from the Sweden Cancerome Analysis Network Breast Initiative is available on GEO (SCAN-B, GSE96058).

Our splicing analysis pipeline v1.0 is available on https://github.com/TheJacksonLaboratory/splicing-pipelines-nf

## Notes

### Competing Interest Statement

The authors have declared no competing interest.

